# Extreme genomic volatility characterises the evolution of the immunoglobulin heavy chain locus in teleost fishes

**DOI:** 10.1101/752063

**Authors:** William J. Bradshaw, Dario Riccardo Valenzano

**Affiliations:** Max Planck Institute for Biology of Ageing, Joseph-Stelzmann-Str. 296, 50937 Cologne, Germany; CECAD Research Center, University of Cologne, Joseph-Stelzmann-Str. 26, 50937 Cologne, Germany

## Abstract

The evolution of the adaptive immune system has provided vertebrates with a uniquely sophisticated immune toolkit, enabling them to mount precise immune responses against a staggeringly diverse range of antigens. Like other vertebrates, teleost fishes possess a complex and functional adaptive immune system; however, our knowledge of the complex antigen-receptor genes underlying its functionality has been restricted to a small number of experimental and agricultural species, preventing a systematic investigation of how these crucial gene loci evolve. Here, we analyse the genomic structure of the immunoglobulin heavy chain (*IGH*) gene loci in the cyprinodontiforms, a diverse and important group of teleosts present in many different habitats across the world. We reconstruct the complete *IGH* loci of the turquoise killifish (*Nothobranchius furzeri*) and the southern platyfish (*Xiphophorus maculatus*) and analyse their *in vivo* gene expression, revealing the presence of species-specific splice isoforms of transmembrane *IGHM*. We further characterise the *IGH* constant regions of ten additional cyprinodontiform species, including guppy, amazon molly, mummichog and mangrove killifish. Phylogenetic analysis of these constant regions reveals multiple independent rounds of duplication and deletion of the teleost-specific antibody class *IGHZ* in the cyprinodontiform lineage, demonstrating the extreme volatility of *IGH* evolution. Focusing on the cyprinodontiforms as a model taxon for comparative evolutionary immunology, this work provides novel genomic resources for studying adaptive immunity and sheds light on the evolutionary history of the adaptive immune system.

## Introduction

The ancient evolutionary arms race between hosts and parasites has given rise to a wide variety of highly sophisticated offensive and defensive adaptations in different taxa^1^. Among the most complex and effective of these adaptations is the vertebrate adaptive immune system, in which developing B- and T-lymphocytes generate a vast diversity of novel antigen-receptor sequences through dynamic recombination of their genomic sequence^1–3^. By combining this enormous diversity in antigen specificities with antigen-dependent clonal expansion and long-term immune memory^4, 5^, vertebrates can progressively improve their protection against recurrent immune challenges while also coping effectively with rapidly-evolving pathogenic threats^6^, dramatically improving their ability to survive and thrive in a complex immune environment.

The immunoglobulin heavy chain (*IGH*) is one of the most important antigen-receptor genes in the adaptive immune system, determining both the effector function and the majority of the antigen-specificity of the antibodies produced by each B-cell^7, 8^. The native structure of the *IGH* gene locus has a profound effect on adaptive immunity in a species, determining the range of gene segment choices available for the VDJ recombination process giving rise to novel antigen-receptor sequences^2^, the possible antibody classes (or *isotypes*) available, and the relationship between VDJ recombination and isotype choice^9^. Understanding the structure of this locus is therefore essential for understanding adaptive-immune function in any given vertebrate species, while comparing loci between species can provide important insight into the adaptive immune system’s complex evolutionary history^9^.

The teleost fishes are the largest and most diverse group of vertebrates, with nearly 30,000 species comprising almost half of extant vertebrate diversity^10^. Previous work has characterised the *IGH* locus structure in a number of teleost species, including zebrafish^11^, medaka^12^, three-spined stickleback^13, 14^, rainbow trout^15^, fugu^16^, and Atlantic salmon^17^. These characterisations have revealed remarkable diversity in the size, structure and functionality of teleost *IGH* loci^9, 18^. However, the number of loci characterised is very small compared to the total evolutionary diversity of teleost fish, and is mainly confined to major aquaculture species and established research models^9, 18^, with characterised species typically quite distantly related to one another within the teleost clade^19^. This relatively sparse sampling of teleost *IGH* loci has prevented higher-resolution analysis of locus structural evolution across groups of closely related species.

Here, we present the first characterisations of *IGH* loci in the Cyprinodontiformes, a large order of teleosts with representatives in diverse habitats and ecological niches across the world. Complete characterisations were performed on the loci of the turquoise killifish (*Nothobranchius furzeri*) and southern platyfish (*Xiphophorus maculatus*), two important model organisms for ecological and evolutionary research^20–23^, while the loci of ten further species (Fig. 1 and Table S2) underwent partial characterisation with a focus on their constant regions. Comparison of these loci revealed dramatic and unexpected differences in *IGH* locus structure and function, including surprising differences in isotype availability and exon usage among different cyprinodontiform species. Phylogenetic analysis showed that the specialised mucosal antibody isotype *IGHZ* has undergone repeated duplication and convergent loss in the course of cyprinodontiform evolution, indicating an unexpected degree of volatility in the evolution of mucosal adaptive immunity. Taken together, this work significantly extends our knowledge of constant-region diversity in teleost fish, and establishes the cyprinodontiforms, and especially the African killifishes, as an ideal model system for comparative evolutionary immunology.

**Figure 1:**
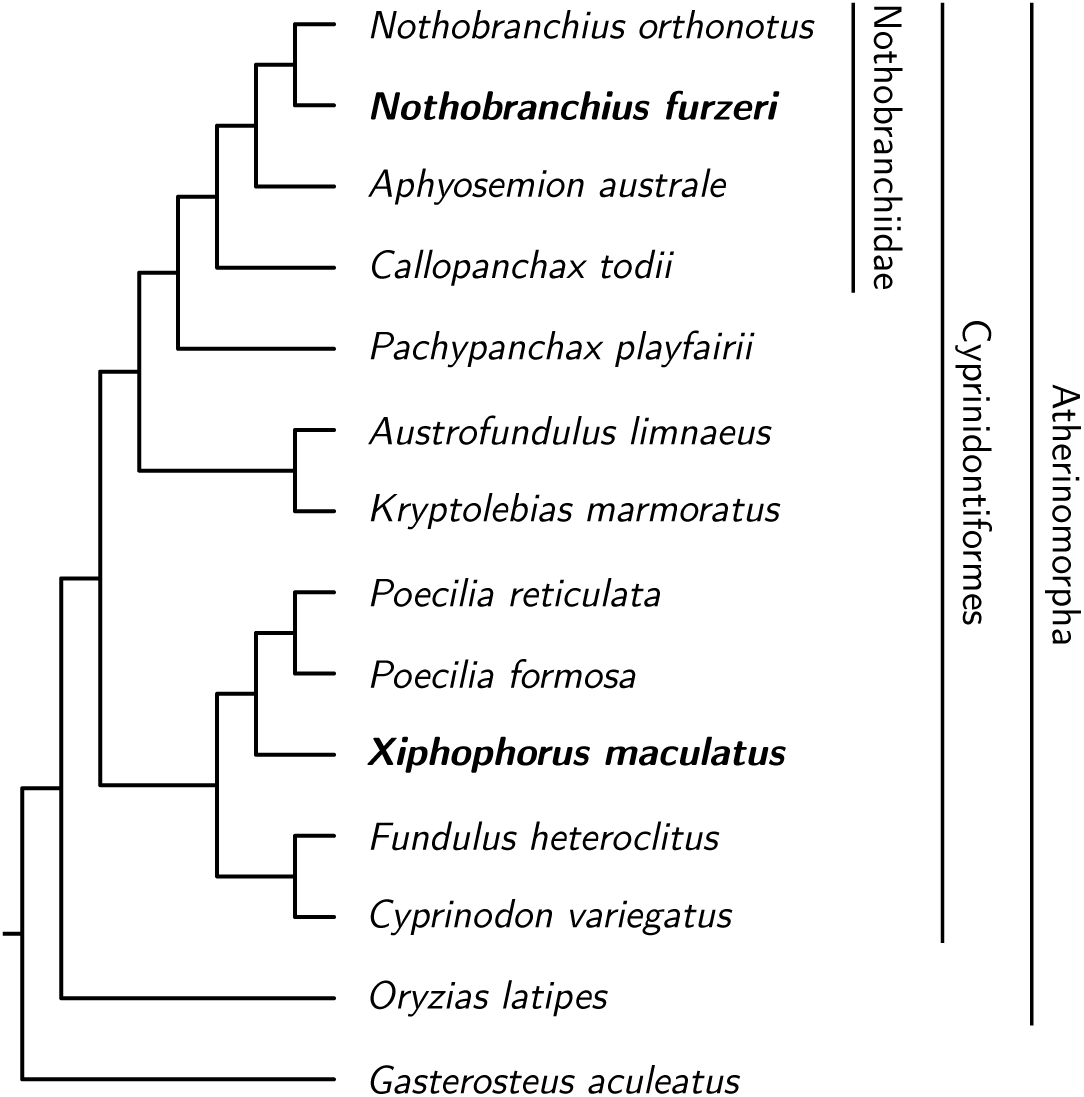
Cladogram of species included in the *IGH* locus analysis. Boldface type indicates species for which new, complete *IGH* locus assemblies were generated for this study; other species were either previously-characterised reference species (*G. aculeatus*, *O. latipes*) or underwent constant-region characterisation only (all other species). Labelled vertical bars designate; higher taxa of interest.

## Results

### The *IGH* loci of *N. furzeri* and *X. maculatus* are highly distinct

In order to assemble and characterise the *IGH* loci in *N. furzeri* and *X. maculatus*, published *IGH* gene segments from zebrafish^11^, medaka^12^ and stickleback^13, 14^ were aligned to the most recent genome assemblies of *N. furzeri* and *X. maculatus* (Table S2) using BLAST^24, 25^. In *X. maculatus*, a single promising region was identified on chromosome 16, while in the *N. furzeri* genome a single region on chromosome 6 and a number of unaligned scaffold sequences were identified as potentially containing parts of the locus. In order to determine which of the candidate scaffolds were genuine parts of the *N. furzeri IGH* locus and integrate them into a continuous locus sequence, bacterial artificial chromosome (BAC) clones from the killifish genomic BAC library^21^ were identified on the basis of alignment of their end sequences to promising genome scaffolds, sequenced on an Illumini MiSeq machine and assembled using SPAdes^26^ and SSPACE^27^, with final refinements made using end-to-end PCR and Sanger sequencing^28^. The resulting BAC inserts were integrated with the identified genome scaffolds (Fig. S3) to produce a single, contiguous locus sequence, on which *IGH* gene segments were identified through more stringent alignment to sequences from reference species (Methods).

The *IGH* locus in *Nothobranchius furzeri* occupies roughly 306 kb on chromosome 16 (NFZ v2.0, accession TBD), while that of *Xiphophorus maculatus* occupies roughly 293 kb on chromosome 16 (scaffold NC_036458.1, Genbank accession GCA_002775205.2). While similar in size, the two loci differ markedly in organisation and content: while the *N. furzeri* locus comprises two distinct subloci on opposite strands (*IGH1* and *IGH2*, Fig. 2a), that of *X. maculatus* forms a single long configuration without any additional subloci (Fig. 2b). The two subloci of the *N. furzeri* locus exhibit a very high degree of synteny with one another in the JH and constant regions, while the VH and DH regions are more divergent, with what appear to be repeated deletion events in the VH/DH regions of *IGH2* (Fig. 2c).

**Figure 2:**
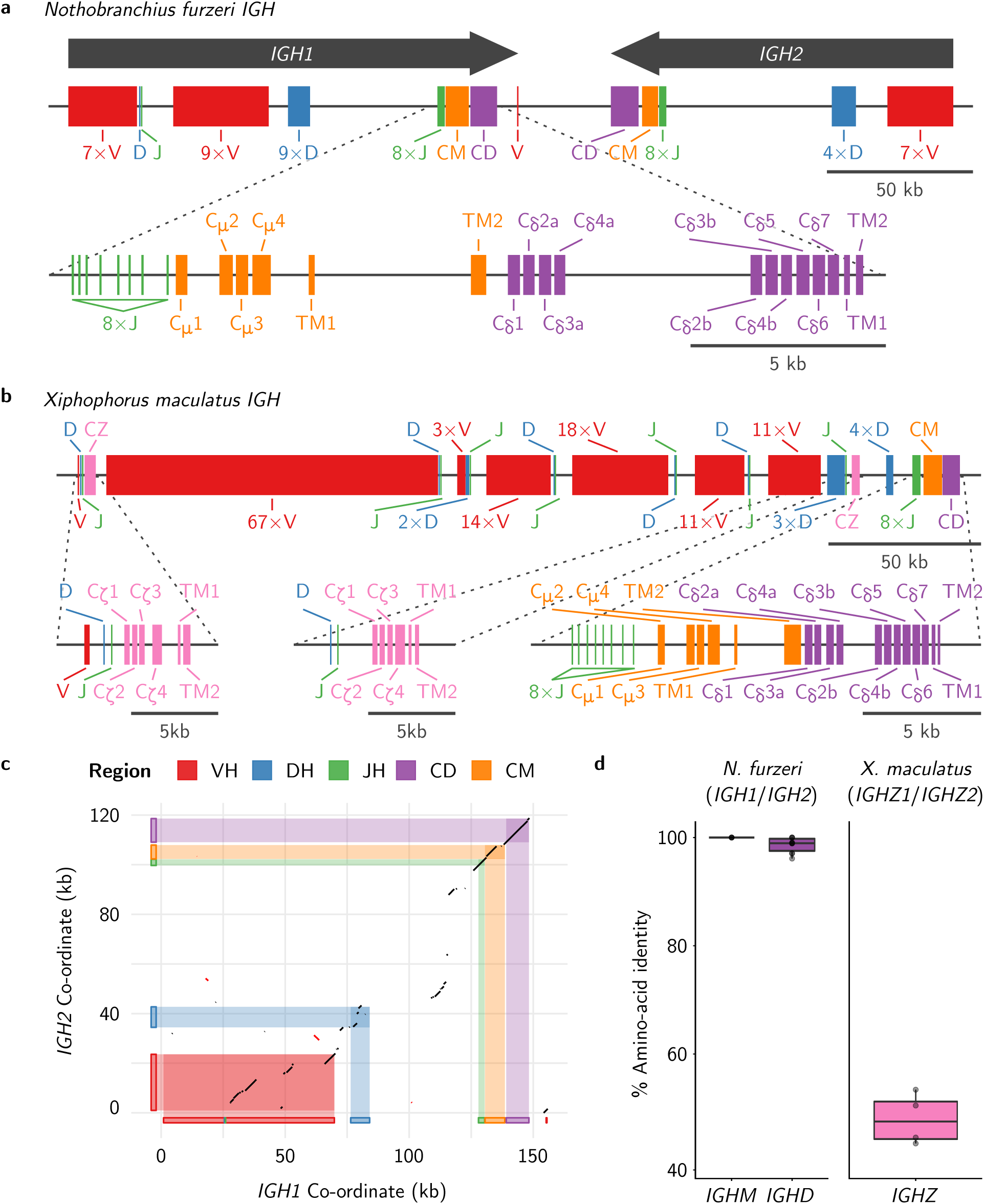
*IGH* locus structure in *Nothobranchius furzeri* and *Xiphophorus maculatus*. **a**, Arrangement of VH, DH, JH and constant regions on the *N. furzeri IGH* locus, indicating the two subloci *IGH1* and *IGH2* and the detailed exon composition of the *IGH1* constant regions. **b**, VH, DH, JH and constant regions on the *X. maculatus IGH* locus, indicating the detailed exon composition of each constant region. **c**, Synteny dot plot of sequential best matches between *N. furzeri IGH1* and *IGH2* sequences, with gene-segment regions in each sublocus indicated by coloured rectangles along each axis. **d**, Boxplots of percentage amino-acid sequence identity between corresponding C*_µ_* and C*_δ_* exons in *N. furzeri IGH1* vs *IGH2* subloci (left) or between corresponding C*_ζ_* exons in *X. maculatus IGHZ1* vs *IGHZ2* constant regions (right).

Three constant-region isotypes have been observed in previously-published teleost loci: *IGHM* and *IGHD*, which are universal in teleosts and homologous to the isotypes of the same names in mammals, and *IGHZ* (also known as *IGHT*), which is teleost-specific and absent in a minority of previously published loci^9, 18^. *X. maculatus IGH*, *N. furzeri IGH1* and *N. furzeri IGH2* all contain intact and highly similar *IGHM* and *IGHD* constant regions, with a six-exon C*_µ_* 1-C*_µ_* 2-C*_µ_* 3-C*_µ_* 4-TM1-TM2 configuration for *IGHM* and a twelve-exon C*_δ_* 1-(C*_δ_* 2-C*_δ_* 3-C*_δ_* 4)_2_-C*_δ_* 5-C*_δ_* 6-C*_δ_* 7-TM1-TM2 configuration for *IGHD* (Fig. 2a and 2b). Such expansion of *IGHD* through tandem duplications of the C*_δ_* 2-C*_δ_* 3-C*_δ_* 4 exons is common in teleosts and has also been observed in zebrafish, channel catfish and Atlantic salmon^9^. Secretory forms of *IGHD* have previously been observed in a minority of teleost loci, produced via either a specialised secretory exon^29^ or a post-C*_δ_* 7 secretory tail^30^; however, neither of these configurations could be found in either *N. furzeri* or *X. maculatus*, and it may be the case that *IGHD* is expressed solely in transmembrane form in these species.

Previous work in rainbow trout has shown that, while *IGHM* is primarily responsible for the serum response to antigenic stimulus, the mucosal response in at least some teleost species is primarily mediated by *IGHZ*^31, 32^, suggesting that this isoform has a specialised mucosal role analogous to *IGHA* in mammals. Unlike *IGHM* and *IGHD*, *IGHZ* is completely absent from both subloci of the *N. furzeri IGH* locus. In contrast, the *X. maculatus IGH* locus contains two distinct *IGHZ* constant regions: *IGHZ1* and *IGHZ2*. *IGHZ2*, like most *IGHZ* constant regions in characterised teleost loci^9^, is located downstream of the VH region and upstream of the larger DH and JH regions preceding *IGHM*; in contrast, and much more unusually, *IGHZ1* is located at the far 5’ end of the *X. maculatus* locus (Fig. 2b). Despite sharing a common six-exon C*_ζ_* 1-C*_ζ_* 2-C*_ζ_* 3-C*_ζ_* 4-TM1-TM2 configuration (Fig. 2b), these two paralogous constant regions are highly distinct, with an average of only 48.0 % amino-acid sequence identity between corresponding C*_ζ_* exons (Fig. 2d), indicating a relatively ancient origin; in contrast, corresponding C*_µ_* and C*_δ_* exons in the two *N. furzeri IGH* subloci exhibit an average of 100 % and 98.6 % amino-acid sequence identity across subloci respectively (Fig. 2d), suggesting a much more recent duplication event.

In terms of the variable regions of the *IGH* gene, the most striking difference between the two loci is in the total number of VH regions: 125 in *X. maculatus* compared to only 24 in *N. furzeri*. In contrast, the number of DH and JH regions are similar between the two species, with 14 DH and 17 JH segments in *N. furzeri* and 14 DH and 15 JH in *X. maculatus*. In *X. maculatus*, only a single VH, DH and JH segment are present upstream of *IGHZ1*, suggesting only a single V/D/J combination is available to antibodies of this isotype; most other segments are present in six V*_n_*D_1_*_−_*_3_J_1_ blocks between *IGHZ1* and *IGHZ2*, with larger blocks of DH and JH segments between *IGHZ2* and *IGHM*. This (V-D-J)*_n_*-C block structure, which is also observed in *N. furzeri IGH1*, is in some ways intermediate between the classic translocon configuration seen in most teleost *IGH* loci and the multi-cluster configuration observed in sharks^18, 33^.

### *N. furzeri* and *X. maculatus* express distinct forms of transmembrane *IGHM*

The six-exon genomic structure of the *IGHM* constant region is highly conserved across the jawed vertebrates, with similar configurations observed in mammals, teleost fishes and elasmobranchs^9, 18^. In all these groups, the choice between secretory and transmembrane *IGHM* is made via alternative splicing following transcription, with the secretory form consistently adopting a four-exon C*_µ_* 1-C*_µ_* 2-C*_µ_* 3-C*_µ_* 4 configuration. Transmembrane *IGHM*, in contrast, differs in configuration between taxa^9^: in mammals, a cryptic splice site within C*_µ_* 4 is used to connect the transmembrane exons, while in teleosts the canonical splice site at the end of C*_µ_* 3 is typically used, excising C*_µ_* 4. Unusually, however, the primary configuration of *IGHM-TM* in medaka (*Oryzias latipes*) has been found to differ from that of other teleosts, with C*_µ_* 2 spliced directly to TM1 and excising C*_µ_* 3 and C*_µ_* 4^9, 12^ (Fig. 3a). Given this surprising diversity, we decided to investigate which splice isoforms are expressed in *N. furzeri* and *X. maculatus*.

**Figure 3:**
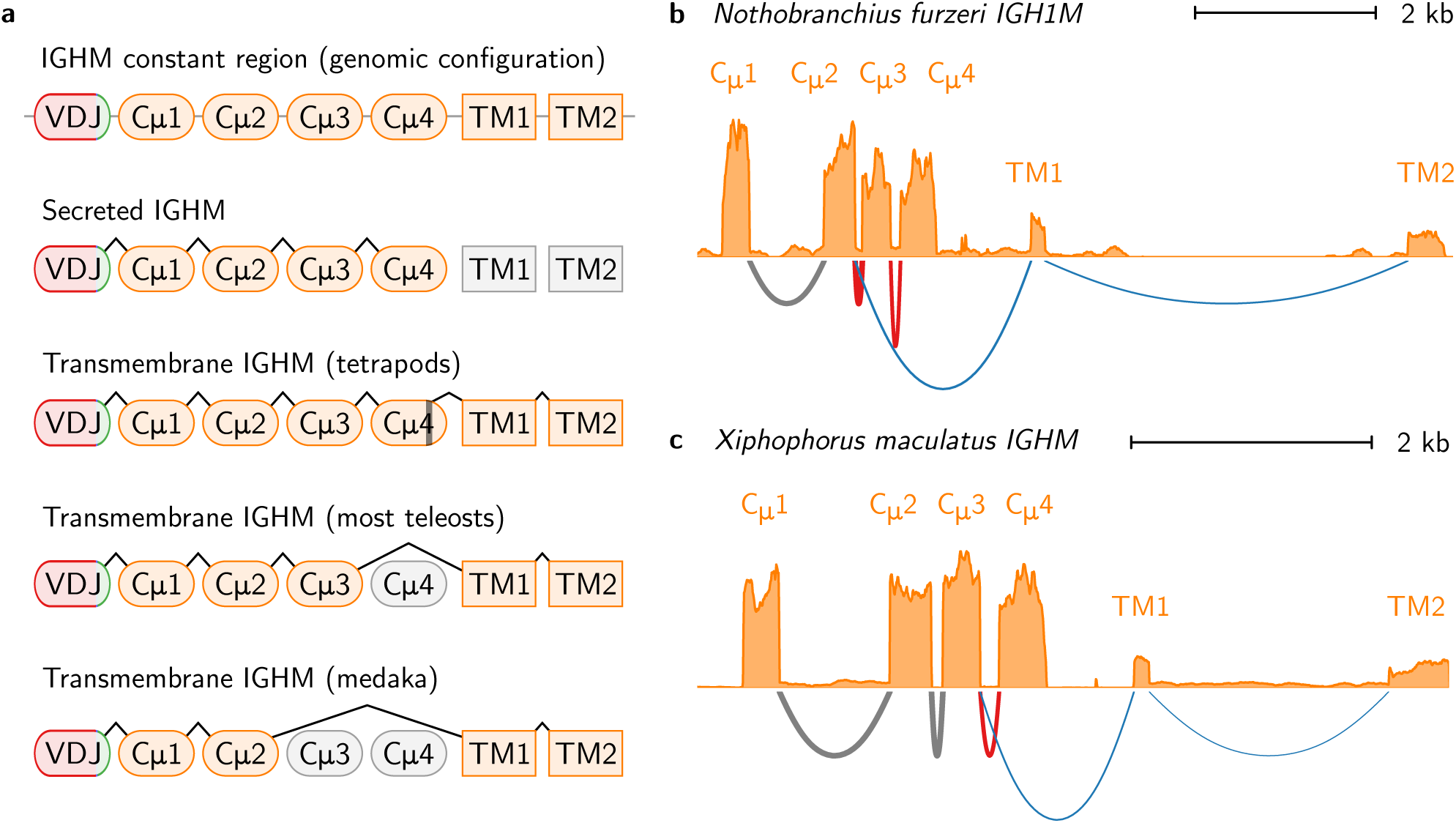
RNA-sequencing data reveals distinct transmembrane isoforms of *IGHM* in *X. maculatus* and *N. furzeri*. **a**, Schematic of *IGHM* splice isoforms in different vertebrate taxa^9^. **b-c**, Read coverage histograms and Sashimi plots of alignment and splicing behaviour of RNA-sequencing reads aligned to the *IGHM* constant regions of **a**, *X. maculatus* and **b**, *N. furzeri*, showing the alternative splicing of transmembrane (blue) and secreted (red) isoforms in both species and the difference in exon usage in *IGHM-TM* between species.

To investigate the exon configuration of expressed *IGH* mRNA in *N. furzeri* and *X. maculatus*, published RNA-sequencing reads from both species (Table S3) were mapped to their respective *IGH* loci using STAR^34^. Surprisingly, the results revealed that the two species utilised different exon configurations for transmembrane *IGHM*: in *X. maculatus*, the standard teleost five-exon configuration was used (Fig. 3c), while *N. furzeri* utilised the unusual four-exon configuration seen in medaka (Fig. 3b), demonstrating that both configurations persist within the cyprinodontiform lineage.

In contrast to *IGHM*, both *N. furzeri* and *X. maculatus* shared a common configuration of transmembrane *IGHD*, with all twelve exons expressed in series. As in other teleosts^9^, expressed *IGHD* in both species began with a chimeric C*_µ_* 1 exon from the upstream *IGHM* constant region (Fig. S1). In *X. maculatus*, meanwhile, both *IGHZ1* and *IGHZ2* expressed a six-exon transmembrane isoform, while *IGHZ1* was also found to give rise to a four-exon secreted isoform comprising C*_ζ_* 1 to C*_ζ_* 4 and a run-on secretory tail; while a tail sequence was also found following C*_ζ_* 4 in *IGHZ2*, no expression of a distinct secretory isoform was detectable in the RNA-sequencing data for this constant region (Fig. S2).

### *IGHZ* has undergone repeated duplication and loss in the Cyprinidontiformes

Medaka (*Oryzias latipes*) is the closest relative of either *N. furzeri* or *X. maculatus* whose *IGH* locus has previously been characterised, and one of the few teleost species previously known to lack the teleost-specific isoform *IGHZ*^9, 12, 18^. Despite this close relationship, the presence of multiple intact *IGHZ* constant regions in *X. maculatus* strongly implies that the absence of this isotype in medaka and *N. furzeri* is the result of two independent deletion events, suggesting that isotype-loss events in teleost *IGH* may be relatively frequent. To investigate this hypothesis in more detail, we identified and characterised *IGH* constant-region sequences in the genomes of ten further cyprinodontiform species (Fig. 1 and Table S2), as well as a new and improved medaka genome assembly (Genbank accession GCA_002234675.1), and investigated the constant-region isoforms present in each species.

The analysed species showed a high degree of variety in locus structure, with dramatic variation in the number and arrangement of constant-region sequences (Fig. 4 and Tables S22 to S24). Of the thirteen species investigated, all had at least one tandem pair of *IGHM* and *IGHD* constant regions, while eight possessed at least one complete *IGHZ* constant region (Fig. 4). Of the exceptions, *Austrofundulus limnaeus* was found to exhibit an orphaned, pseudogenised *IGHZ-TM1* exon but no C*_ζ_* exons in the current genome assembly, while no *IGHZ* exons at all were found in the genomes of *O. latipes*, *N. furzeri*, *Aphyosemion australe*, or *Nothobranchius orthonotus*. Assuming that *IGHZ*, once deleted, cannot be restored to the *IGH* locus in a lineage, a simple visualisation on a species tree (Fig. 5a) confirms that that medaka and *N. furzeri* represent two distinct *IGHZ* deletion events; *A. limnaeus* appears to represent another independent deletion event, for a total of at least three *IGHZ* deletions within the clade containing the cyprinodontiforms and medaka.

**Figure 4:**
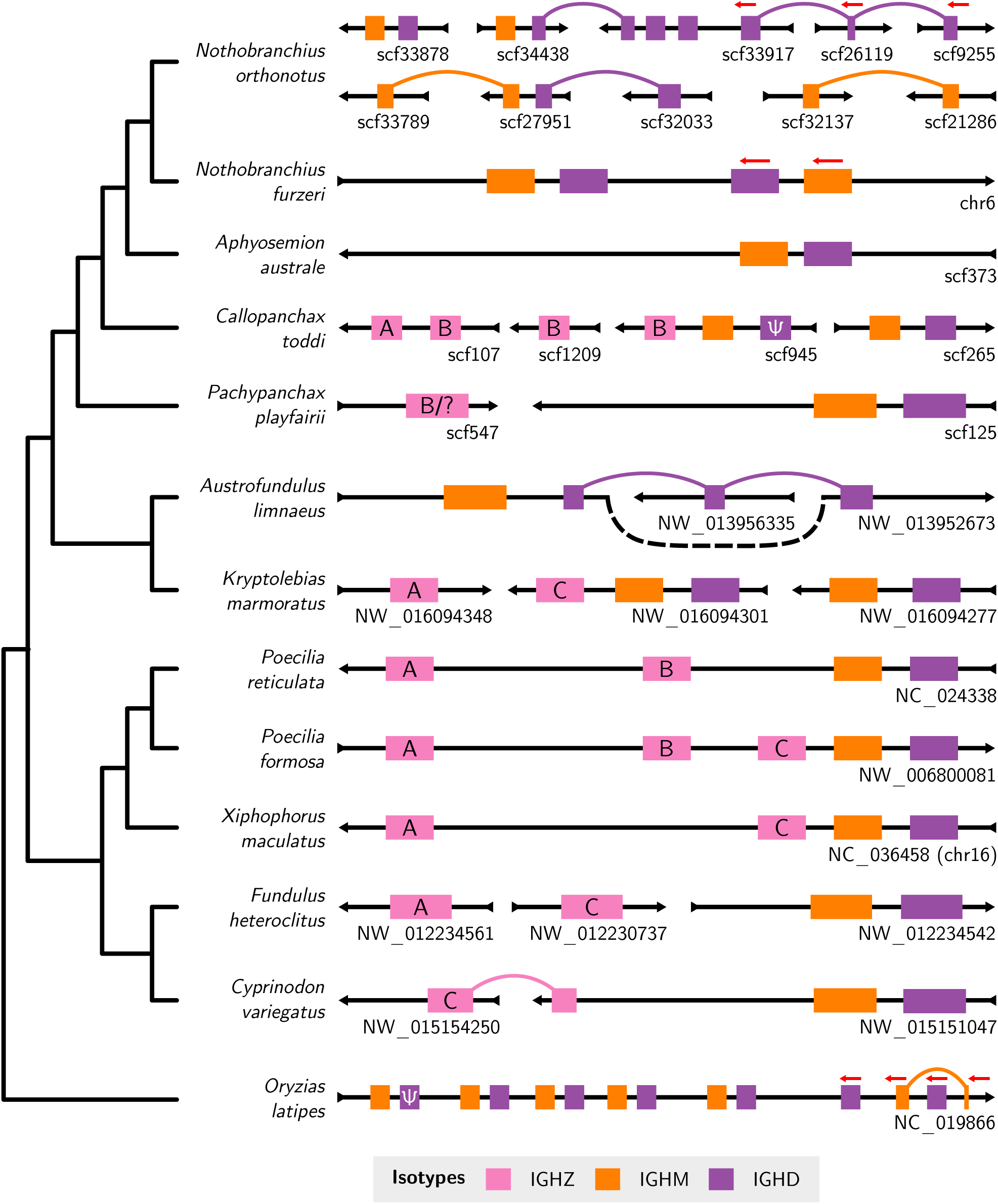
Constant-region organisation in the Atherinomorpha. Schematic of newly-characterised *IGH* constant regions in the genomes of thirteen species from the Atherinomorpha (Cyprinodontiformes + medaka). Scaffold orientation is given by the black arrows; constant regions are oriented left-to-right unless otherwise specified (red arrows). Scaffold names are displayed beneath each scaffold on the right-hand side. Links between regions on different scaffolds indicate that exons from what appears to be the same constant region are distributed across multiple scaffolds in the order indicated; the order of unlinked scaffolds is arbitrary. The isotype of each region is given by its colour; *IGHZ* regions are further annotated with their subclass (Fig. 5b). Clearly pseudogenised constant regions are indicated by Ψ. Isotype length, scaffold length, and scaffold position are not to scale. Variable regions and lone, isolated constant-region exons are not shown. The cladogram to the left indicates evolutionary relationships between species (Fig. 1).

**Figure 5:**
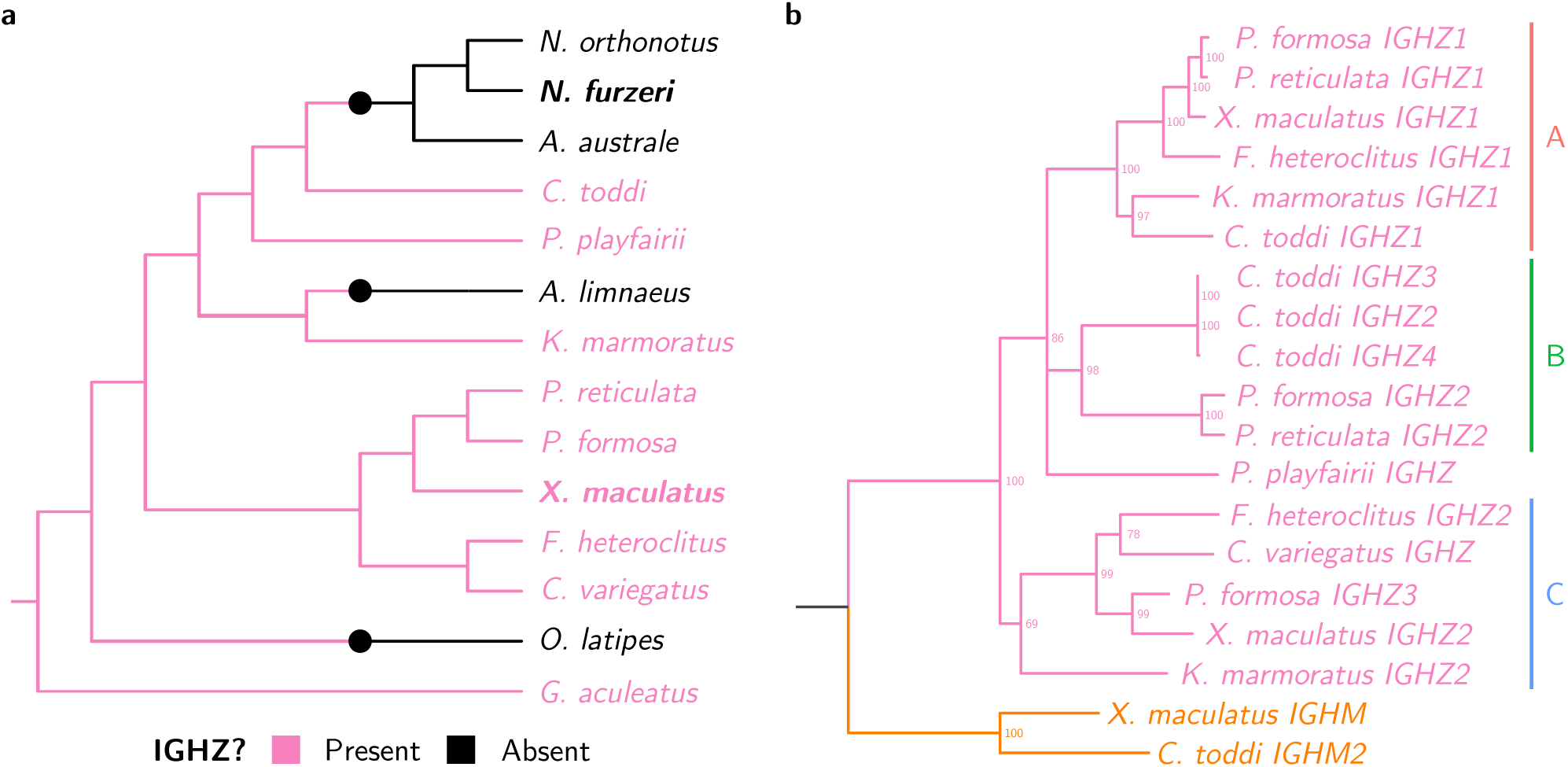
*IGHZ* has undergone repeated duplication and loss in the Cyprinodontiformes. **a**, Cladogram of species from Fig. 1, with three-spined stickleback (*Gasterosteus aculeatus*) as the outgroup, coloured according to known *IGHZ* status. Large coloured points indicate inferred state-change events. **b**, Phylogram of concatenated C*_ζ_* 1-4 nucleotide sequences from *n IGHZ*-bearing Cyprinodontiform species, with C*_µ_* 1-4 sequences from two species as outgroup (in orange). Nodes with less than 65 % bootstrap support are collapsed into polytomies, while major monophyletic subclasses are annotated on the right.

In addition to being lost repeatedly, *IGHZ* also demonstrates a relatively high level of multiplicity within the cyprinodontiforms, with a geometric mean of 1.93 *IGHZ* constant regions per *IGHZ*-bearing locus (a 1.62:1 ratio relative to *IGHM* or *IGHD*). This multiplicity suggests a more complex evolutionary history than can be captured by a simple presence/absence metric. Concordantly, phylogenetic analysis with PRANK^35^ and RAxML^36^ (Fig. 5b, alignment length 1733 bp, 35% gaps/missing characters) reveals three distinct monophyletic clades (or subclasses) of *IGHZ* constant regions in the Cyprinidontiformes, *IGHZA* to *C*, each of which is present in multiple different species and appears to have been present in the common ancestor of the eight IGHZ-bearing species analysed. The only locus whose *IGHZ* could not be assigned to one of these subclasses, that of *Pachypanchax playfairii*, appears to have undergone a fusion event, with *P. playfairii* C*_ζ_* 1 and C*_ζ_* 2 aligning strongly to *IGHZB* exons from other species while *P. playfairii* C*_ζ_* 3 and C*_ζ_* 4 show more ambiguous alignment behaviour favouring *IGHZA* or *IGHZC* (Fig. 6).

**Figure 6:**
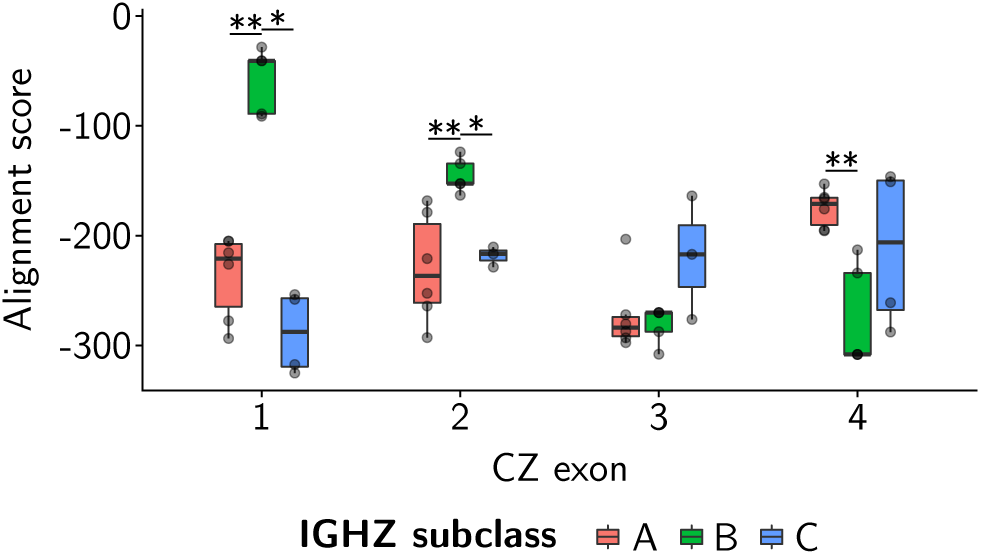
*Pachypanchax playfairii IGHZ* is composed of exons from multiple ancestral subclasses. Box-plots of Needleman-Wunsch alignment scores between the amino-acid sequences of *Pachypanchax playfairii* C*_ζ_* exons and those of equivalent exons from seven other *IGHZ*-bearing cyprinodontiform species, demonstrating the differing affinity of different *P. playfairii* exons for each of the three *IGHZ* subclasses. Less negative scores indicate a stronger alignment. Pairwise *p*-values were computed using nonparametric Mann-Whitney *U* tests (*∗*:0.01 *< p ≤* 0.05; *∗∗*:0.001 *< p ≤* 0.01).

In summary, in addition to the still-universal primitive antibody classes *IGHM* and *IGHD*, the cyprinodontiforms ancestrally possessed at least three subclasses of *IGHZ*, which subsequently evolved in parallel across the clade. Each of these subclasses has been lost in multiple cyprinodontiform species, with different species showing distinct patterns of retention and loss, and in at least one lineage – that of *Pachypanchax playfairii* – two different *IGHZ* lineages appear to have fused to produce a chimeric isotype. All three subclasses are missing from a subset of species in the Nothobranchiidae (including *Nothobranchius furzeri*), and also appear to have been independently lost in *Austrofundulus limnaeus*, further demonstrating the remarkable volatility of the *IGH* locus across evolutionary time.

## Discussion

The immunoglobulin heavy chain locus is notable for its size and complexity, as well as for the central role it plays in vertebrate adaptive immunity and survival. Previous research in teleost fishes has revealed a remarkable degree of diversity in the length, organisation, and isotype composition of different *IGH* loci^9, 18^, with important but understudied implications for antibody diversity and immune functionality among teleost species.

In this study, we presented the first detailed characterisations of *IGH* loci from the Cyprinodontiformes, a widespread order of teleost fishes that include many important model systems in evolutionary biology and ecology. Two such species, the turquoise killifish *Nothobranchius furzeri* and the southern platyfish *Xiphophorus maculatus*, underwent complete assembly and characterisation of their *IGH* loci, while ten other cyprinodontiform species received partial characterisations focused on their constant regions. These additional species were selected on the basis of their relatedness to *N. furzeri* and *X. maculatus* and their prevalence in the research literature, and included a number of prominent ecological model organisms (including guppy^37^, mummichog^38^ and mangrove rivulus^39^), yielding a dataset with significant relevance to researchers studying the role of infection and immunity in teleost ecology.

The *IGH* loci of *X. maculatus* and *N. furzeri* exhibited radically different locus organisations, with dramatic differences in VDJ number, locus organisation and isotype availability. These results are consistent with previous findings of highly-diverse teleost loci and support a process of rapid locus evolution in the cyprinodontiforms. Characterisation of the constant regions of additional cyprinodontiform species confirmed this finding, with several groups of closely-related species (e.g. *Nothobranchius furzeri*, *Nothobranchius orthonotus* and *Callopanchax toddi*) showing highly divergent locus structures and constant-region availability (Fig. 4).

It is interesting to speculate on the origins of this extremely rapid diversification in gene structure. Very little is known about the relationship between environmental context and immune locus structure; it is possible that part of the variety in *IGH* gene locus structure in the Cyprinodontiformes represents divergent adaptations to different immune environments. Alternatively, this diversification may be primarily the result of unusually high rates of stochastic, non-adaptive changes in gene structure in germline *IGH*, or to relaxation of selective constraints on locus structure. Finally, at least some of the difference between locus structures in different species is likely to be attributable to differences in assembly quality; for example, the characterisation of medaka constant regions presented here contains many fewer unusual or incomplete constant regions than that presented in the published medaka *IGH* locus^12^, primarily due to the increased quality of the more recent medaka genome assemblies. Issues with assembly quality could also account for the apparent complexity of the *Nothobranchius orthonotus* locus, as the genome of this species was assembled from a wild-caught individual with a high degree of heterozygosity^40^.

The teleost-specific isotype *IGHZ* is widespread among teleost species, and appears to play a specialised role in mucosal immunity^31, 32^. Before the publication of this work, only two teleost species (medaka and channel catfish) were known or thought to lack the *IGHZ* antibody isotype in their *IGH* loci, suggesting that the loss of *IGHZ* may be a relatively rare event. However, in addition to confirming the absence of *IGHZ* in medaka, the work presented here has identified four new teleost species (*Nothobranchius furzeri*, *Nothobranchius orthonotus*, *Aphyosemion australe* and *Austrofundulus limnaeus*) that appear to lack *IGHZ* constant regions in their *IGH* loci, representing two distinct and previously unknown loss events independent from that affecting the closely-related medaka. This finding, which triples the number of known teleost species without *IGHZ* and doubles the number of known loss events, is even more striking when combined with the discovery that the cyprinidontiform common ancestor likely had no fewer than three distinct *IGHZ* constant regions (Fig. 5b), all of which would have to be lost on the way to any *IGHZ*-free lineage. Taken together, these observations suggest that the presence/absence of *IGHZ* in the wider teleost clade may be much more volatile than suggested by previously available locus data, and raises the possibility that, given sufficiently high-density analysis of other teleost lineages, a surprisingly high frequency of *IGHZ*-lacking species may also be found elsewhere.

The absence of *IGHZ* from so many species in this analysis naturally raises the important question of how the mucosal adaptive immune system in these species differs from that of their *IGHZ*-bearing relatives: how, and to what extent, can the primitive isotype *IGHM* compensate for the loss of a specialised mucosal antibody class? This question is especially interesting in the case of *IGHZ*-lacking species with close *IGHZ*-bearing relatives (e.g. *Nothobranchius furzeri* and *Callopanchax toddi*, or *Austrofundulus limnaeus* and *Kryptolebias marmoratus*); if it is the case that mucosal immune responses differ systematically between these species, such that *IGHM* takes up some or all of the roles normally played by *IGHZ*, then uncovering the mechanisms by which this shift is regulated could reveal important new insights into decision-making and control of humoral adaptive immunity. Similarly, characterising the different functional roles and responses of different *IGHZ* subclasses in cyprinodontiform fishes could yield important information about how these species interact with different aspects of their immune environment.

Another important difference between *N. furzeri* and *X. maculatus*, whose evolution is more difficult to investigate using genomic data, is the exon-usage behaviour of expressed *IGHM*. In *X. maculatus*, transmembrane *IGHM* adopts the same configuration as that seen in most teleosts: a five-exon isoform in which the end of C*_µ_* 3 is spliced to the start of TM1 and C*_µ_* 4 is excised. Conversely, in *N. furzeri IGHM-TM* adopts the same four-exon configuration observed in medaka, in which C*_µ_* 3 is also excluded. Given that *X. maculatus* adopts the primitive configuration, the recurrence of the same unusual configuration in both medaka and turquoise killifish is surprising, and indicates that both configurations are present in the Cyprinodontiformes; more information about the evolutionary history of this divergence in splicing behaviour, along with data on the functional consequences of including or excluding C*_µ_* 3 from the transmembrane protein structure of IGHM, could yield important new insights into antibody evolution and functionality in teleost fishes.

One of the most important advances in immunology in recent years has been the explosion of quantitative, high-throughput approaches for investigating the composition, diversity and functionality of the antibody repertoire^41–43^. As a direct result of the research presented here, twelve previously-uncharacterised teleost species now have databases of *IGH* constant-region sequences available, enabling these immunoglobulin-sequencing approaches to be applied in the cyprinodontiforms for the first time. Combining antibody-repertoire data with other information gathered from wild fishes could yield important new insights into the role of the adaptive immune system in the lives and evolution of wild vertebrates. In addition, the possibility of sequencing the repertoires of several related species adds an exciting comparative dimension previously missing in immune-repertoire studies, opening up the possibility of simultaneously comparing the response of different closely-related species to a common immunogenic stimulus. This comparative element would be especially interesting in the context of investigating the repertoire responses of closely related species with different *IGHZ* genotypes, as well as for comparing the functional roles of different *IGHZ* subclasses across species.

In combination with the genomic and functional findings discussed above, such large-scale comparative repertoire studies provide a novel opportunity for comparative evolutionary immunology in the Cyprinodontiformes, with the potential to greatly expand our knowledge of the interaction between ecological conditions and the evolution of the adaptive immune system in teleost fishes.

## Methods

### Assembling the Nothobranchius furzeri IGH locus

To identify promising candidate sequences from which to assemble the *N. furzeri IGH* locus sequence, VH, JH and CH sequences from three reference species with published *IGH* loci (zebrafish^11^, medaka^12^ and three-spined stickleback^13, 14^) were aligned to the most recent assembly of the *N. furzeri* genome^44^ (NFZ v2.0, Accession TBD) using BLAST^24, 25^. Scaffolds containing promising alignments to at least two distinct types of *IGH* gene segment, or which covered at least 1 % of the total length of the scaffold, were retained as potentially containing parts of the *IGH* locus.

In order to determine which of these candidate scaffolds contained parts of the *IGH* locus and integrate them into a single sequence, clones from the killifish genomic BAC library^21^ were identified on the basis of alignment of their end sequences to promising genome scaffolds. These BAC clones were provided to us by the FLI in Jena and isolated and sequenced as described in the next section.

Following sequencing, demultiplexed and adapter-trimmed MiSeq reads were processed with Trimmotatic^45^ to trim low quality sequence and Bowtie 2^46^ to remove contaminating *E. coli* sequences, then corrected with QuorUM^47^ or BayesHammer^26, 48^ and assembled with SPAdes^26^. Following assembly, any *E. coli* scaffolds resulting from residual contaminating reads were identified by aligning scaffolds to the *E. coli* genome using BLASTN^24, 25^, and scaffolds containing significant matches were discarded. The remaining scaffolds were then scaffolded using SSPACE^27^ using jumping libraries from the killifish genome project^20, 21, 44^.

In order to guarantee the reliability of the assembled scaffolds, the assemblies produced with BayesHammer- and QuorUM-corrected reads were compared, and scaffolds were broken into segments whose contiguity was agreed on between both assemblies. To integrate these fragments into a contiguous insert assembly, points of agreement between BAC assemblies from the same genomic region (e.g. two scaffolds from one assembly aligning concordantly to one scaffold from another) and between BAC assemblies and genome scaffolds, were used to combine scaffolds where possible. Any still-unconnected scaffolds were assembled together through pairwise end-to-end PCR using Kapa HiFi HotStart ReadyMix PCR Kit according to the manufacturer’s instructions, followed by Sanger sequencing^28^ (Eurofins). PCR primers for end-to-end PCR were designed using Primer3^49^.

Following BAC insert assembly, assembled inserts were screened for *IGH* locus segments in the same manner described for genome scaffolds above. Passing BAC inserts were aligned to the candidate genome scaffolds and chromosome sequence with BLASTN and integrated manually (Fig. S3), giving priority in the event of a sequence conflict to (i) any sequence containing a gene segment missing from the other, and (ii) the genome scaffold sequence if neither sequence contained such a segment. BACs and scaffolds which could not be integrated into the locus sequence in this way were discarded as orphons.

### BAC isolation and sequencing

All BAC clones that were sequenced for this research were provided by the FLI in Jena as plate or stab cultures of transformed *E. coli*, which were replated and stored at 4*^◦^*C. Prior to isolation, the clones of interest were cultured overnight in at least 100 ml LB medium. The resulting liquid cultures were transferred to 50 ml conical tubes and centrifuged (10-25 min, 4*^◦^*C, 3500*g*) to pellet the cells. The supernatant was carefully discarded and the cells were resuspended in 18 ml QIAGEN buffer P1.

After resuspension, the cultures underwent alkaline lysis to release the BAC DNA and precipitate genomic DNA and cellular debris. 18 ml QIAGEN buffer P2 was added to each tube, which was then mixed gently but thoroughly by inversion and incubated at room temperature for 5 min. 18 ml ice-chilled QIAGEN neutralisation buffer P3 was added to precipitate genomic DNA and cellular debris, and each tube was mixed gently but thoroughly by inversion and incubated on ice for 15 min. The tubes were then centrifuged (20-30 min, 4*^◦^*C, 12000*g*) to pellet cellular debris and the supernatant was transferred to new conical tubes. This process was repeated at least two more times, until no more debris was visible in any tube; this repeated pelleting was necessary to minimise contamination in each sample, as the normal column- or paper-based filtering steps used during alkaline lysis resulted in the loss of the BAC DNA.

Following alkaline lysis, the DNA in each sample underwent isopropanol precipitation: 0.6 volumes of room-temperature isopropanol were added to the clean supernatant in each tube, followed by 0.1 volumes of 3 mol sodium acetate solution. Each tube was mixed well by inversion, incubated for 10-15 min at room temperature, then centrifuged (30 min, 4*^◦^*C, 12000*g*) to pellet the DNA. The supernatant was discarded and the resulting DNA smear was “resuspended” in 1 ml 100 % ethanol and transferred to a 1.5 ml tube, which was re-centrifuged (5 min, 4*^◦^*C, top speed) to obtain a concentrated pellet. Finally, the pelleted samples were resuspended in QIAGEN buffer EB and purified of proteins and RNA using standard phenol-chloroform extraction and ethanol precipitation techniques.

The resuspended BAC isolates were sent to the Cologne Center for Genomics, where they underwent Illumina Nextera XT library preparation and were sequenced on an Illumina MiSeq sequencing machine (MiSeq Reagent Kit v3, 2*×*300 bp reads).

### Identifying locus scaffolds in other species

Candidate *IGH* locus sequences in other species (Table S2) were identified in the same manner as for *N. furzeri*, by aligning VH, JH and CH sequences from reference species to available genome sequences with BLAST. In the case of *X. maculatus* the reference species used were zebrafish, stickleback, medaka and *N. furzeri*, while for all other species the gene segments from the *X. maculatus* locus were also used. Additional sequence refinement with BAC inserts was not necessary in these species: in the case of *X. maculatus* only a single sequence region (on chromosome 16) was identified, while in the other species a complete locus characterisation (requiring a single contiguous sequence) was not performed.

### Characterising constant-region sequences and expression

Constant-region sequences on candidate locus scaffolds (or, in the case of *N. furzeri* and *X. maculatus*, on complete locus sequences) were identified by mapping CH sequences from reference species to candidate sequences using BLAST. Following alignment of reference sequences, overlapping alignments to reference segments of the same isotype and exon number were collapsed together, keeping track of the number of collapsed alignments and the best E-values and bitscores obtained for each alignment group. Alignment groups with a very poor maximum E-value (*>* 0.001) were discarded, as were groups overlapping with a much better alignment to a different isotype or exon type, where “much better” was here defined as a bitscore difference of at least 16.5 Where conflicting alignments to different isotypes or exon types co-occurred without a sufficiently large difference in bitscore, both alignment groups were retained for manual resolution of exon identity.

Following resolution of conflicts, alignment groups underwent a second filtering step of increased stringency, requiring a minimum E-value of 10*^−^*^8^ and at least two aligned reference exons over all reference species to be retained. Each surviving alignment group was then converted to a sequence range, extended by 10 bp at each end to account for truncated alignments failing to cover the ends of the exon, and used to extract the corresponding exon sequence into FASTA format. These sequences then underwent manual curation to resolve conflicting exon identities, assign exon names and perform initial end refinement based on putative splice junctions (Tables S4 and S11).

In order to validate intron/exon boundaries and investigate splicing behaviour among *IGH* constant-region exons in *N. furzeri* and *X. maculatus*, published RNA-sequencing data (Table S3) were aligned to the annotated locus using STAR^34^. In both cases, reads files from multiple individuals were concatenated and aligned together, and the *IGH* locus was masked using RepeatMasker^50^ (using the built-in zebrafish repeat parameters) prior to mapping. Mapped reads spanning predicted exons of more than 10 kb were excluded from the alignment, as were read pairs mapping more than 10 kb apart. Following alignment, the resulting SAM files were processed into sorted, indexed BAM files using SAMtools^51^ and visualised with Integrated Genomics Viewer (IGV^52, 53^) to determine intron/exon boundaries of predicted exons, as well as the major splice isoforms present in each dataset. Read-coverage and Sashimi plots (Fig. 3, S1 and S2) were generated from the alignment data using Gviz^54^.

For species other than *N. furzeri* or *X. maculatus*, intron/exon boundaries were predicted manually based on BLASTN and BLASTP alignments to closely-related species and the presence of conserved splice-site motifs (AG at the 5’ end of the intron, GT at the 3’ end^55^). In cases where no 3’ splice site was expected to be present (e.g. for CM4 or TM2 exons), the nucleotide exon sequence was terminated at the first canonical polyadenylation site (AATAAA if present, otherwise one of ATTAAA, AGTAAA or TATAAA^56^), while the amino-acid sequence was terminated at the first stop codon. In many cases, it was not possible to locate a TM2 exon due to its very short conserved coding sequence (typically only 2 to 4 amino-acid residues^11, 13^).

### Characterising variable-region sequences

Variable-region gene segments in the *N. furzeri* and *X. maculatus* were identified and characterised using different methods depending on segment type. For VH and JH segments, segments from reference species were used to construct a multiple-sequence alignment with PRANK^35^, which was then used by NHMMER^57^ to perform a Hidden-Markov-Model-based search for matching sequences in the locus. The resulting sequence candidates were extended on either end to account for boundary errors, then refined manually. In the case of VH sequences, 3’ ends were identified by the start of the RSS heptamer sequence (consensus CACAGTG^58^), if present, while 5’ ends and FR/CDR boundaries were identified using IMGT/DomainGapAlign^59^ with the default settings; where necessary, IMGT/DomainGapAlign was also used to IMGT-gap the VH segments in accordance with the IMGT unique numbering^60^. For JH segments, 5’ ends were identified using the RSS heptamer sequence, while the 3’ end was identified using the conserved splice-junction motif GTA.

Following extraction and manual curation, VH segments were grouped into families based on their pairwise sequence identity. In order to assign segments to families, the nucleotide sequence of each VH segment in a locus was aligned to every other segment using Needleman-Wunsch global alignment^61^ as implemented in the Biostrings R package^62^, and the resulting matrix of pairwise sequence identities was used to perform single-linkage hierarchical clustering on the VH segments. The resulting dendrogram was cut at 80 % sequence identity to obtain VH families (Fig. S4 to S6). These families were then numbered based on the order of the first-occurring VH segment from that family in the first *IGH* sublocus in which the family is represented, and each VH segment was named based on its parent sublocus, its family, and its order among elements of that family in that sublocus (Table S5 and Tables S12 to S16). JH segments, meanwhile, were named based on their order within their parent sublocus and, in *X. maculatus*, on whether they were upstream of *IGHZ* or *IGHM* constant regions (Tables S9 and S20).

Unlike VH and JH gene segments, DH segments are too short and unstructured to be found effectively using an HMM-based search strategy. Instead, DH segments in assembled loci were located using their distinctive pattern of flanking recombination signal sequences in opposite sense^3^. Potential matches to this pattern were searched for using EMBOSS FUZZNUC^63^, with a high mismatch tolerance (up to 8 mismatches across the whole pattern) to account for deviations from the conserved sequence in either or both of the RSSs. Promising candidate sequences from this search were oriented based on the orientation of flanking VH or JH sequences on the same scaffold, then underwent a second, more stringent filtering step in which sequences lacking the most conserved positions in each RSS (in particular, the initial CA motif in the heptamer sequence^58^) were discarded. Finally, the identified DH candidates were checked manually, candidates without good RSS sequences were discarded, and flanking RSS sequences were trimmed to obtain the DH segment sequences themselves. As with the JH segments, these were numbered based on their order within their parent sublocus and, in the case of *X. maculatus*, on whether they were upstream of *IGHZ* or *IGHM* constant regions (Tables S6 and S18).

### Phylogenetic inference

Cladograms of teleost species (Fig. 1 and 5a) were constructed using phylogenetic information from Cui *et al.*^40^ (for African killifishes) and Hughes *et al.*^19^ (for other species) and visualised using the ggtree R package^64^.

To construct a phylogram of *IGHZ* sequences (Fig. 5b), the nucleotide sequences of C*_ζ_* 1-4 exons from each *IGHZ* constant region in Tables S22 to S24 were concatenated together into a single sequence per constant region and aligned to one another using PRANK^35^. The resulting multiple-sequence alignment was then used to perform maximum-likelihood phylogenetic inference with RAxML^36^, using the SSE3-enabled parallelised version of the software, the standard GTR-Gamma nucleotide substitution model, and built-in rapid bootstrapping with 1000 bootstrap replicates; during tree inference, the third codon position was partitioned into a separate model. The bootstrap-annotated RAxML_bipartitions file was inspected and rooted manually in Figtree^65^ and again visualised using ggtree; during tree visualisation, nodes with bootstrap support of less than 65 % were collapsed into polytomies.

### Inter- and intralocus sequence comparison

Synteny between subloci in the *N. furzeri* locus (Fig. 2c) was analysed using the standard synteny pipeline from the DECIPHER R package^66^, which searches for chains of exact *k*-mer matches within two sequences.

Comparison between constant-region exons, either within the same locus (Fig. 2d) or between loci (Fig. 6) were performed using Needleman-Wunsch exhaustive global alignments^61^, as implemented in the Biostrings R package^62^, using the default scoring parameters from that package.

## Acknowledgements

We would like to thank Kathrin Reichwald for providing the BAC clones used in this study; Mario Ventura and Nicola Lorusso for early help and support with BAC isolation; Bérénice Benayoun, Anton Korobeynikov, Jorge Boucas, Franziska Metge and Bernd Wozny for help and advice with the BAC sequence assembly process; and David Willemsen and Rongfeng Cui for critically reading and reviewing the manuscript. This work was funded by the Max Planck Institute for Biology of Ageing, the Cologne Graduate School of Ageing Research, the Max Planck Society and the DFG Collaborative Research Center 1310.

---

**Figure S1:**
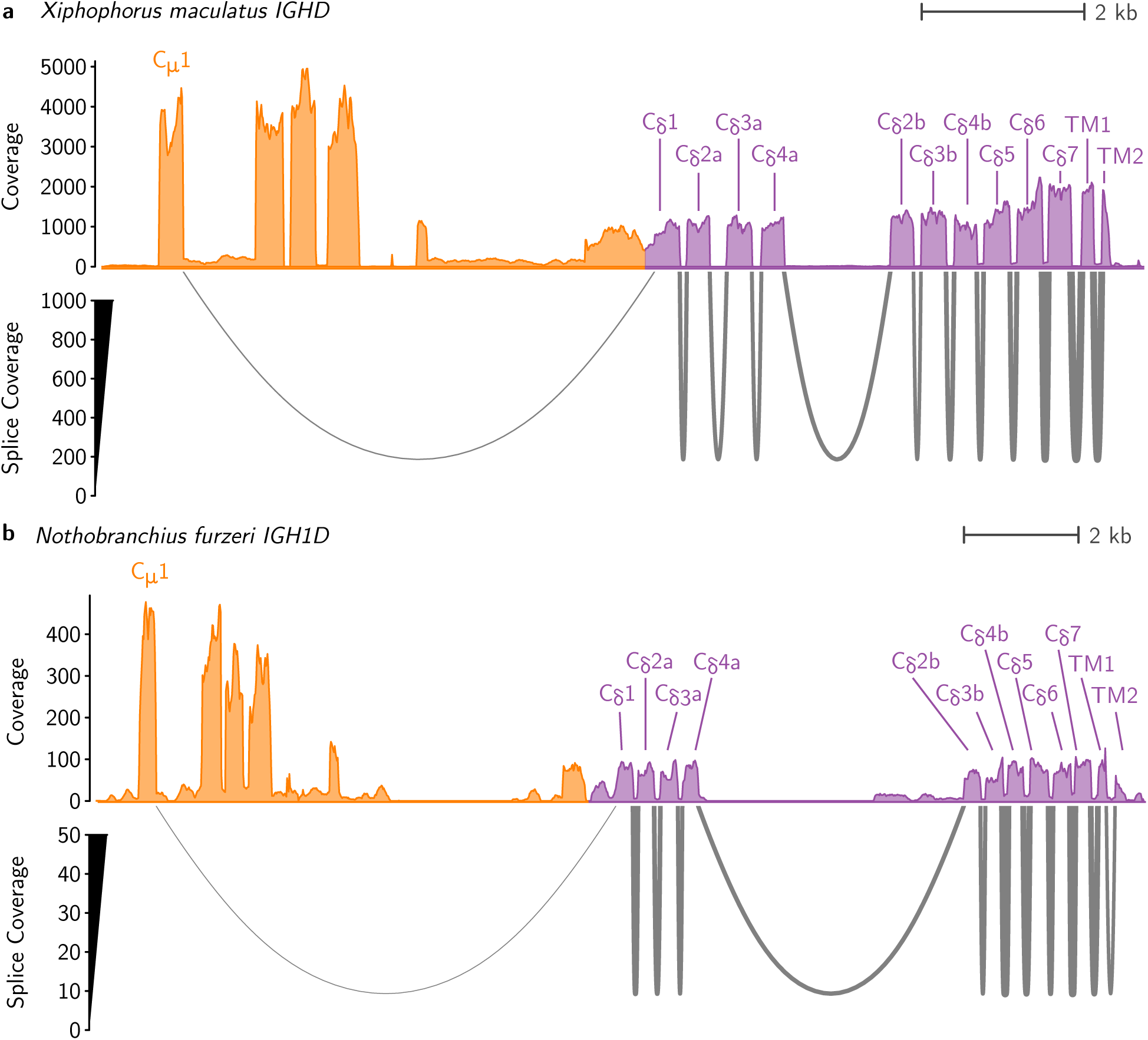
Read coverage and Sashimi plots showing alignment and splicing behaviour of RNA sequencing reads aligned to the *IGHD* constant regions of **a**, *Xiphophorus maculatus* and **b**, *Nothobranchius furzeri*, showing the chimeric splicing of C*_µ_* 1 to the start of the *IGHD* constant region in both species.

**Figure S2:**
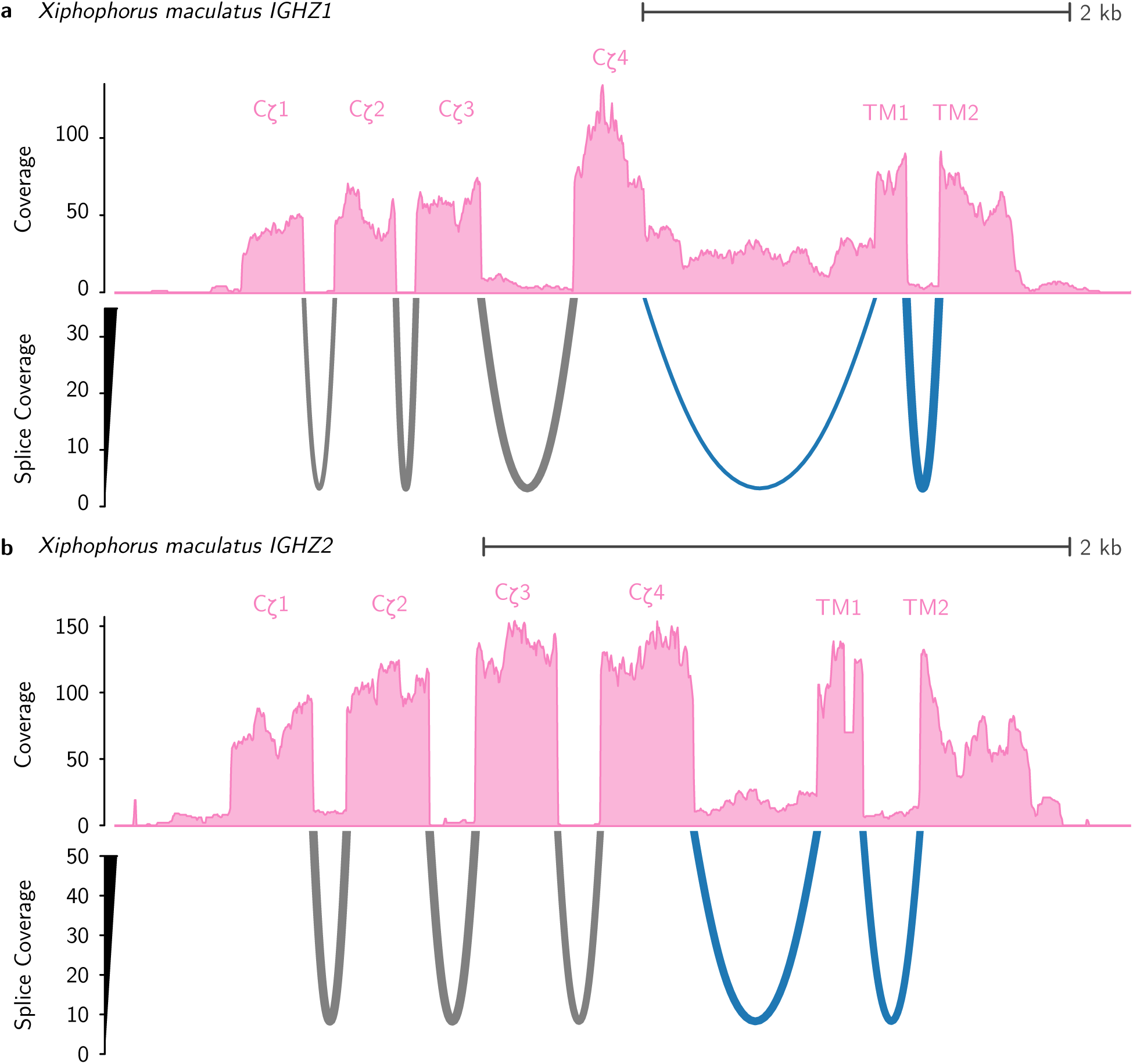
Read coverage and Sashimi plots showing alignment and splicing behaviour of RNA sequencing reads aligned to the (a) *IGHZ1* and (b) *IGHZ2* constant regions of *Xiphophorus maculatus*, showing the alternative splicing of secreted (grey) and transmembrane (grey+blue) isoforms in both cases. Note the apparent expression of a post-splice-site secretory tail after C*_ζ_* 4 in *IGHZ1* but not *IGHZ2*.

**Figure S3:**
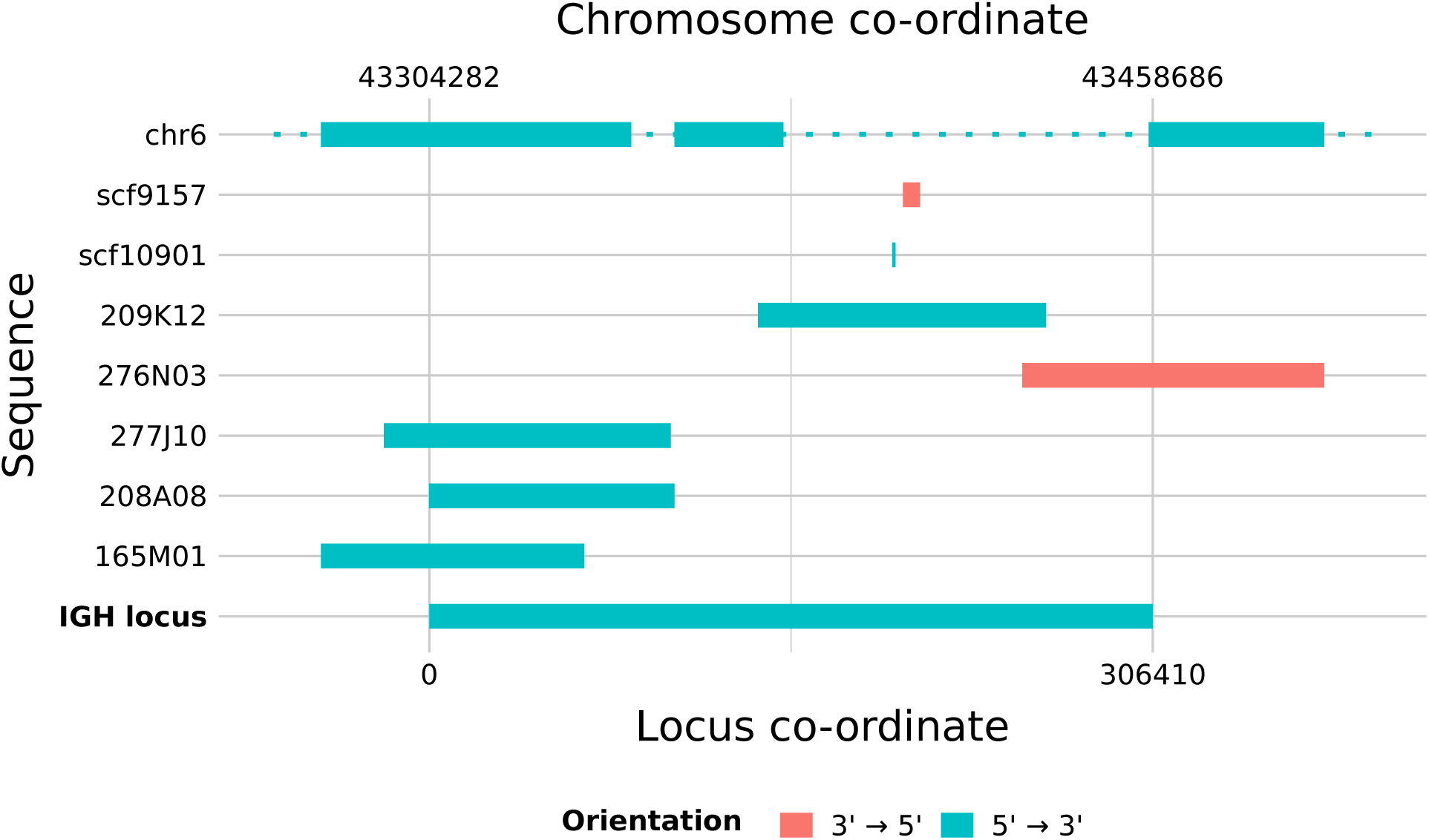
Assembling the *Nothobranchius furzeri IGH* locus: Schematic of genome scaffolds and BAC inserts contributing to the *Nothobranchius furzeri IGH* locus sequence, with their corresponding place within the locus sequence (bottom axis). Internal gaps with dotted lines indicate regions on chromosome 16 with no corresponding locus sequence, as a result of intercalation of BAC or scaffold sequences.

**Figure S4:**
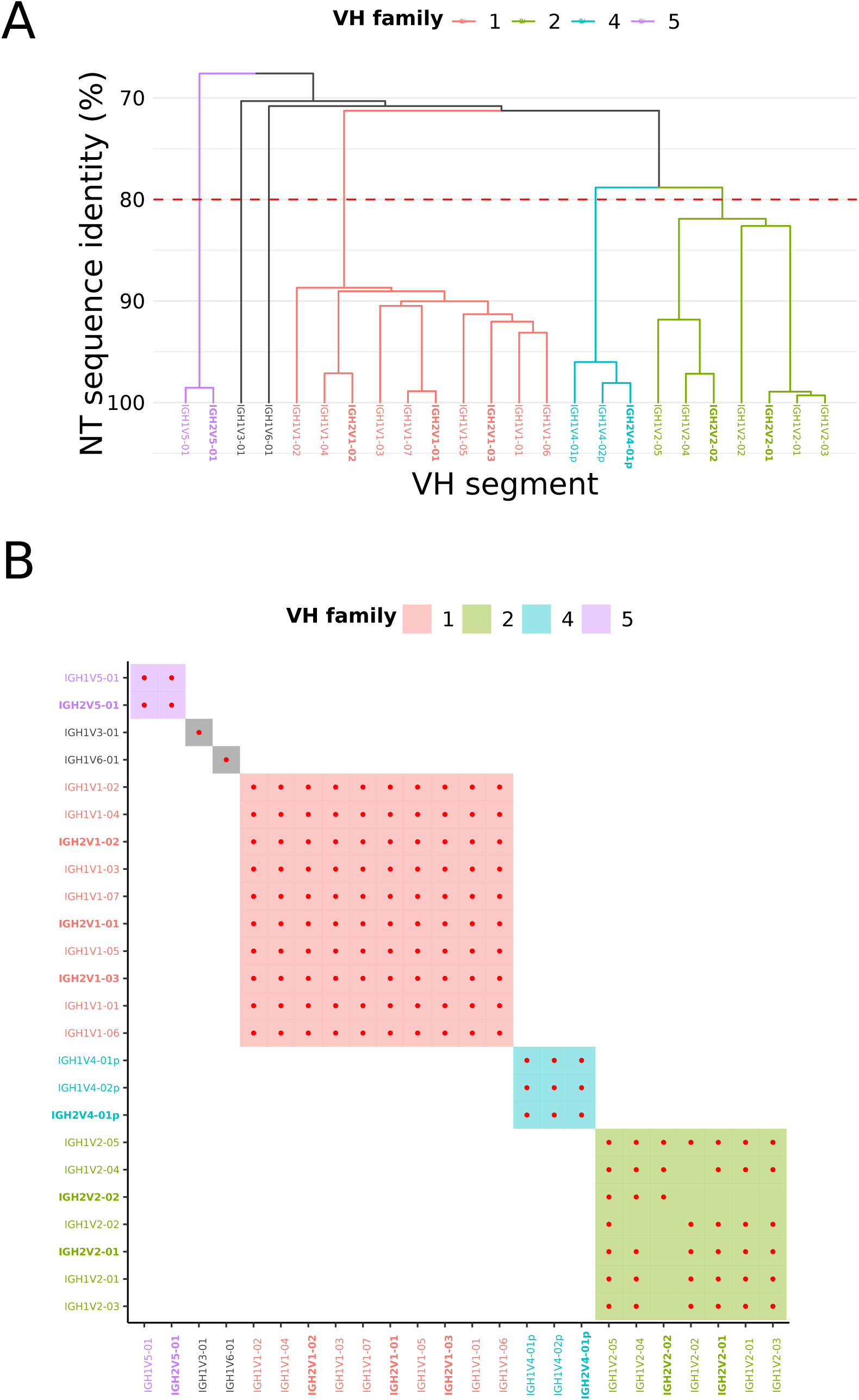
VH families in the *Nothobranchius furzeri IGH* locus: (A) Dendrogram of sequence similarity of VH segments in the *Nothobranchius furzeri IGH* locus, arranged by single-linkage clustering on nucleotide sequence identity. The red line indicates the 80 % cutoff point for family assignment. (B) Heatmap of family relationships among *Nothobranchius furzeri* VH segments, with shaded squares indicating families and red dots indicating pairwise nucleotide sequence identity of at least 80 %. In both subfigures, VH families containing multiple segments are uniquely coloured, single-segment families are in grey, and segments from the *IGH2* sublocus are displayed in boldface.

**Figure S5:**
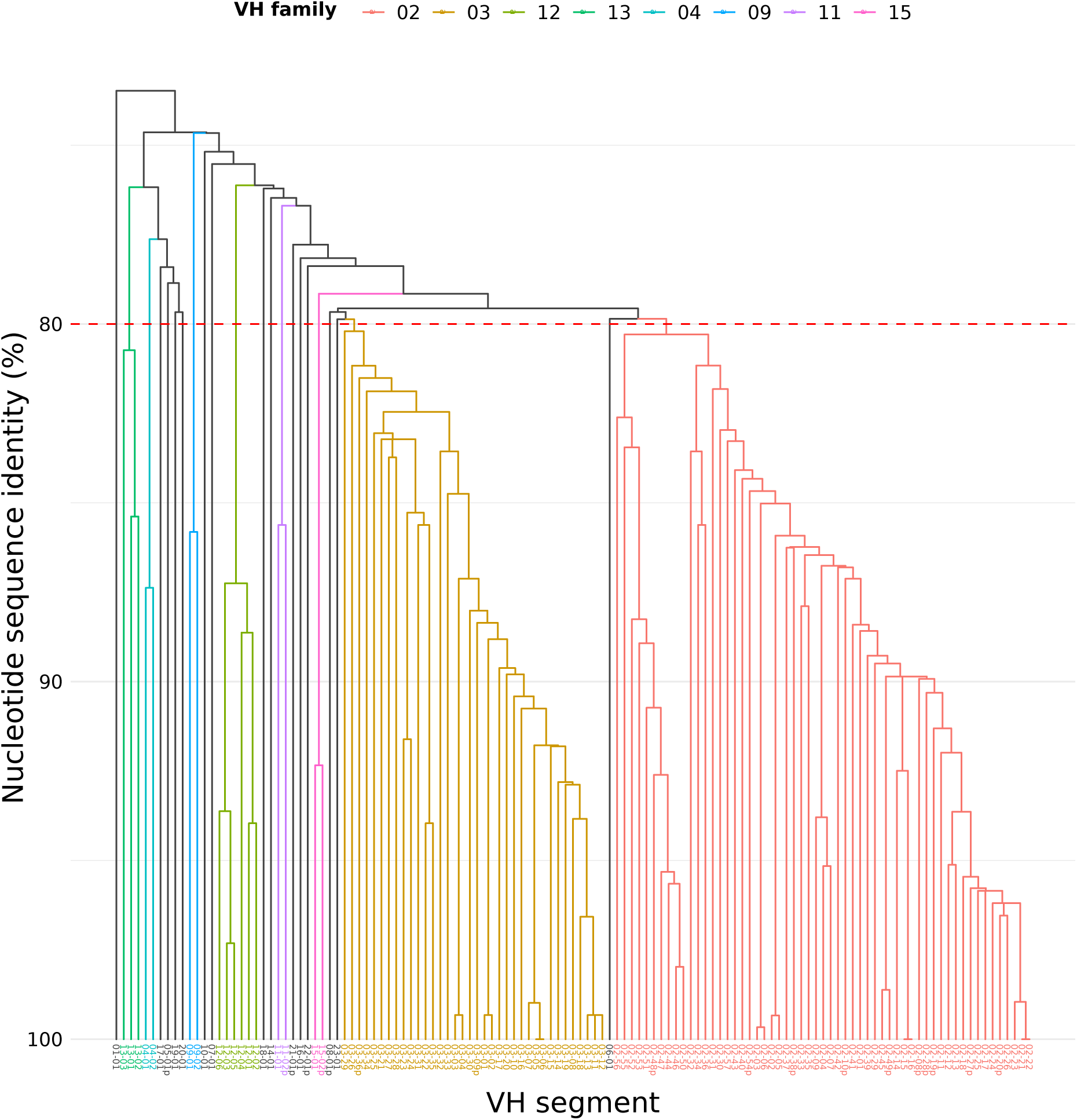
Dendrogram of VH families in the *Xiphophorus maculatus IGH* locus: Dendrogram of sequence similarity of VH segments in the *Xiphophorus maculatus* locus, arranged by single-linkage clustering on nucleotide sequence identity. The red line indicates the 80 % cutoff point for family assignment, while branch colour indicates family membership: VH families containing multiple segments are uniquely coloured, while single-segment families are in grey.

**Figure S6:**
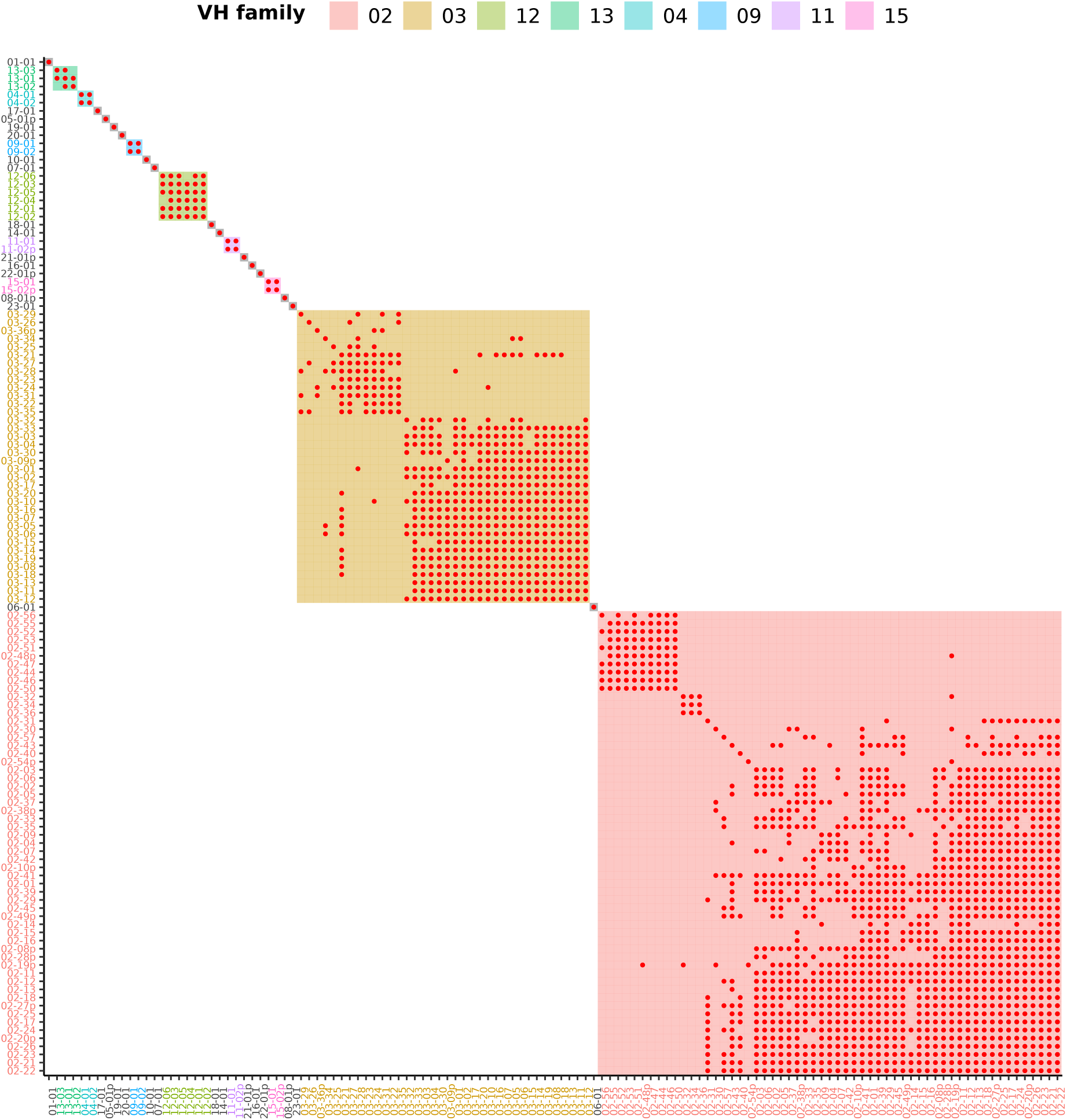
Heatmap of VH families in the *Xiphophorus maculatus IGH* locus: Heatmap of family relationships among *Xiphophorus maculatus* VH segments, with coloured shading indicating families and red dots indicating pairwise nucleotide sequence identity of at least 80%. VH families containing multiple segments are uniquely coloured, while single-segment families are in grey.

**Figure S7:**
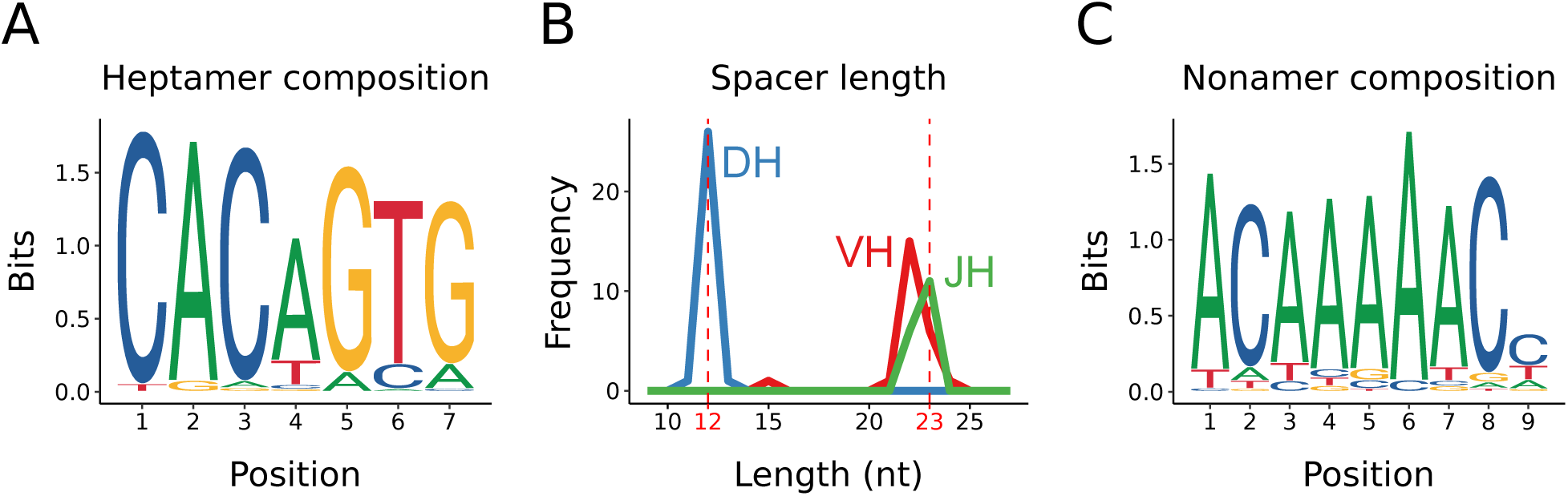
Recombination signal sequences in *Nothobranchius furzeri IGH*: (A) Sequence composition of conserved heptamer sequences across all *Nothobranchius furzeri* heavy-chain RSSs; (B) length distribution of unconserved spacer sequences in *Nothobranchius furzeri* heavy-chain RSSs; (C) sequence composition of conserved heptamer sequences across all *Nothobranchius furzeri* heavy-chain RSSs.

**Figure S8:**
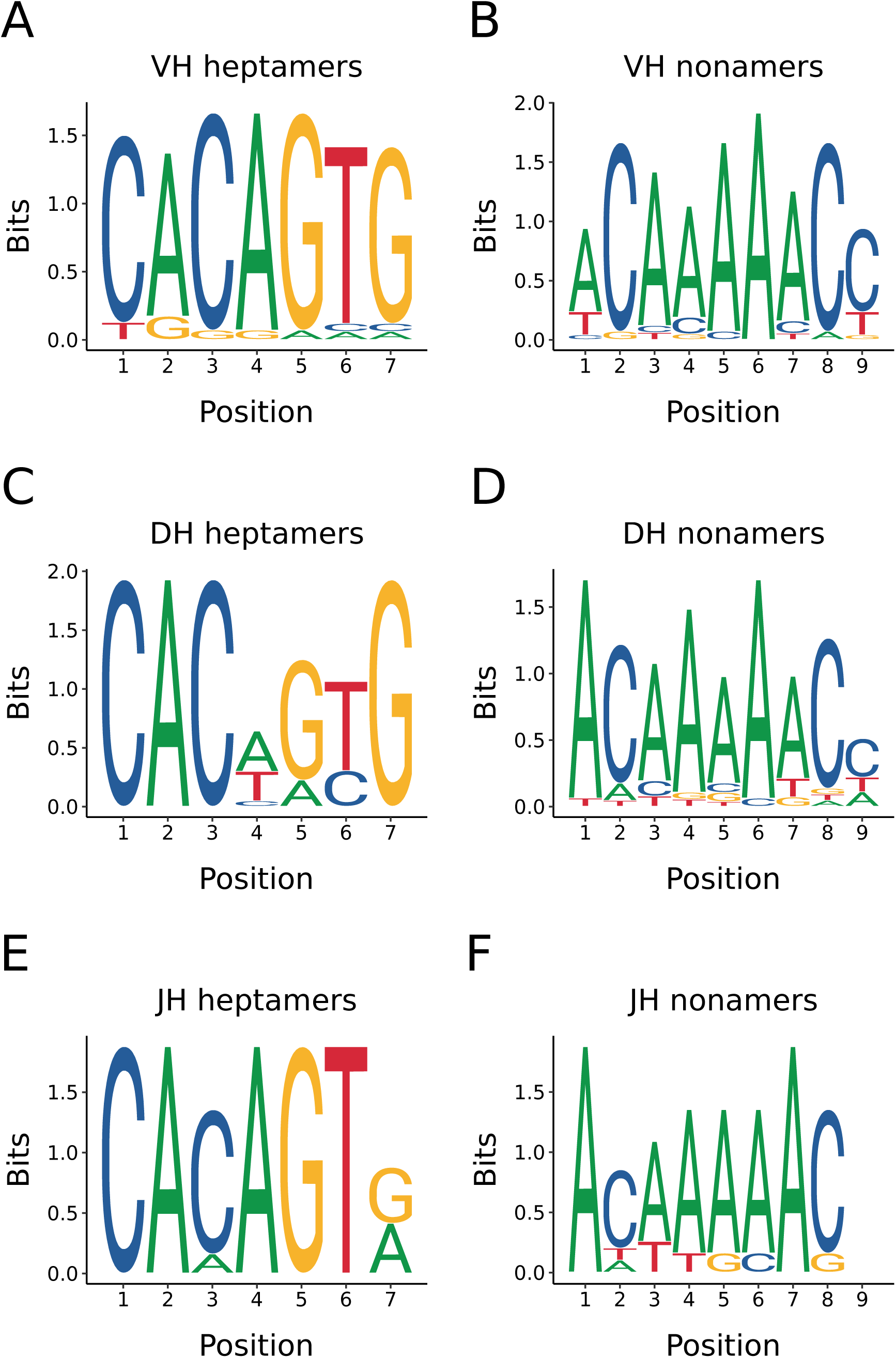
N*o*thobranchius *furzeri* recombination signal sequences by segment type: Sequence composition of conserved heptamer (A,C,E) and nonamer (B,D,F) sequences from *Nothobranchius furzeri* heavy-chain RSSs associated with VH (A,B), DH (C,D) or JH (E,F) gene segments.

**Figure S9:**
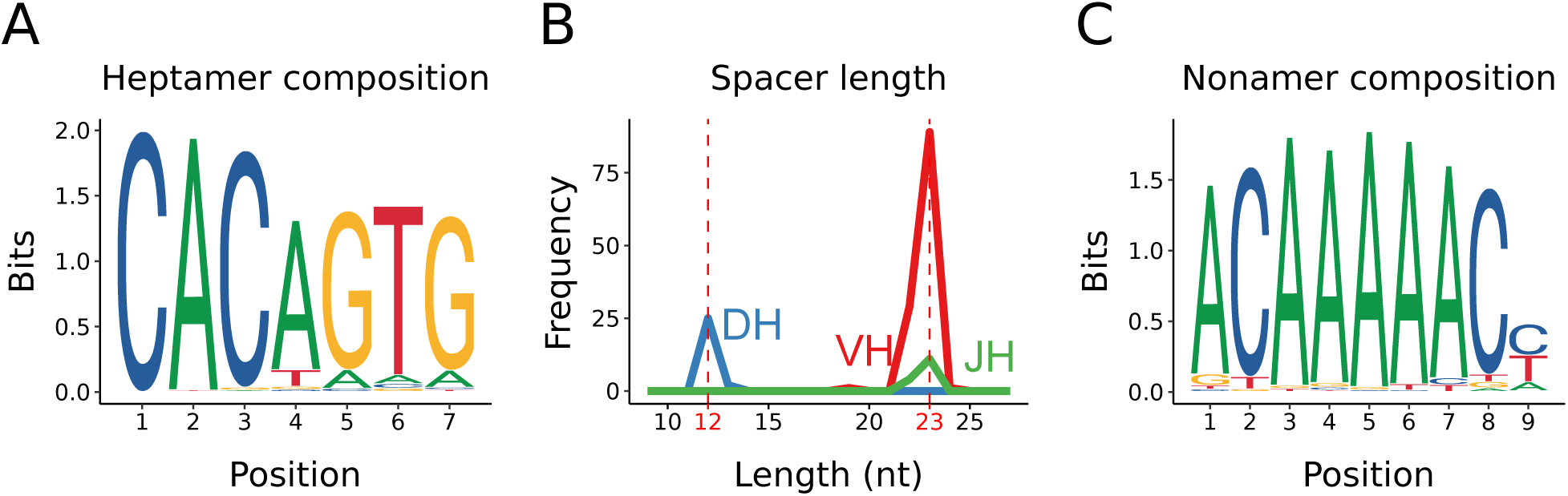
Recombination signal sequences in the *Xiphophorus maculatus IGH* locus: (A) Sequence composition of conserved heptamer sequences across all *Xiphophorus maculatus* heavy-chain RSSs; (B) length distribution of unconserved spacer sequences in *Xiphophorus maculatus* heavy-chain RSSs; (C) sequence composition of conserved heptamer sequences across all *Xiphophorus maculatus* heavy-chain RSSs.

**Figure S10:**
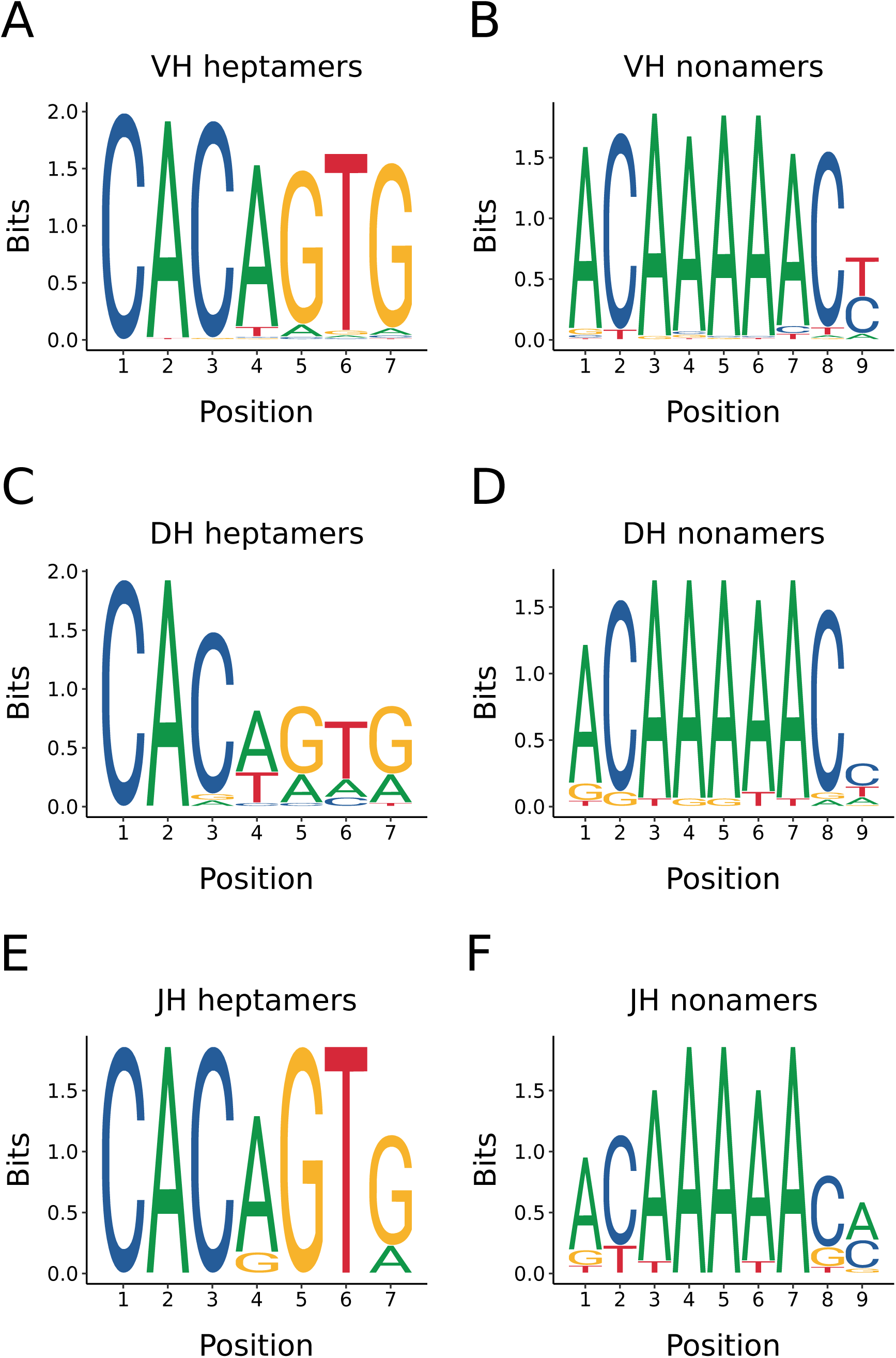
X*i*phophorus *maculatus* recombination signal sequences by segment type: Sequence composition of conserved heptamer (A,C,E) and nonamer (B,D,F) sequences from *X. maculatus* heavy-chain RSSs associated with VH (A,B), DH (C,D) or JH (E,F) gene segments.

**Table S1:**
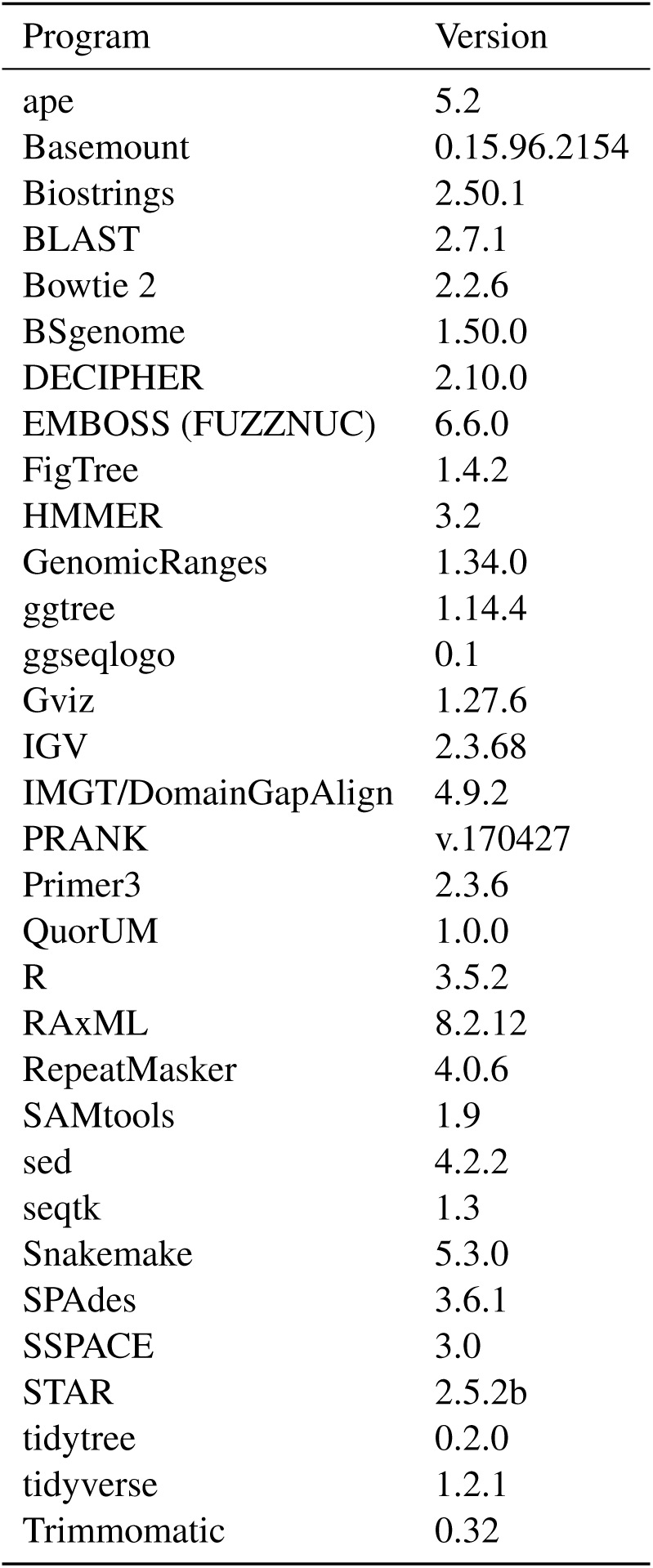
Versions of software and R packages used in computational analyses

**Table S2:**
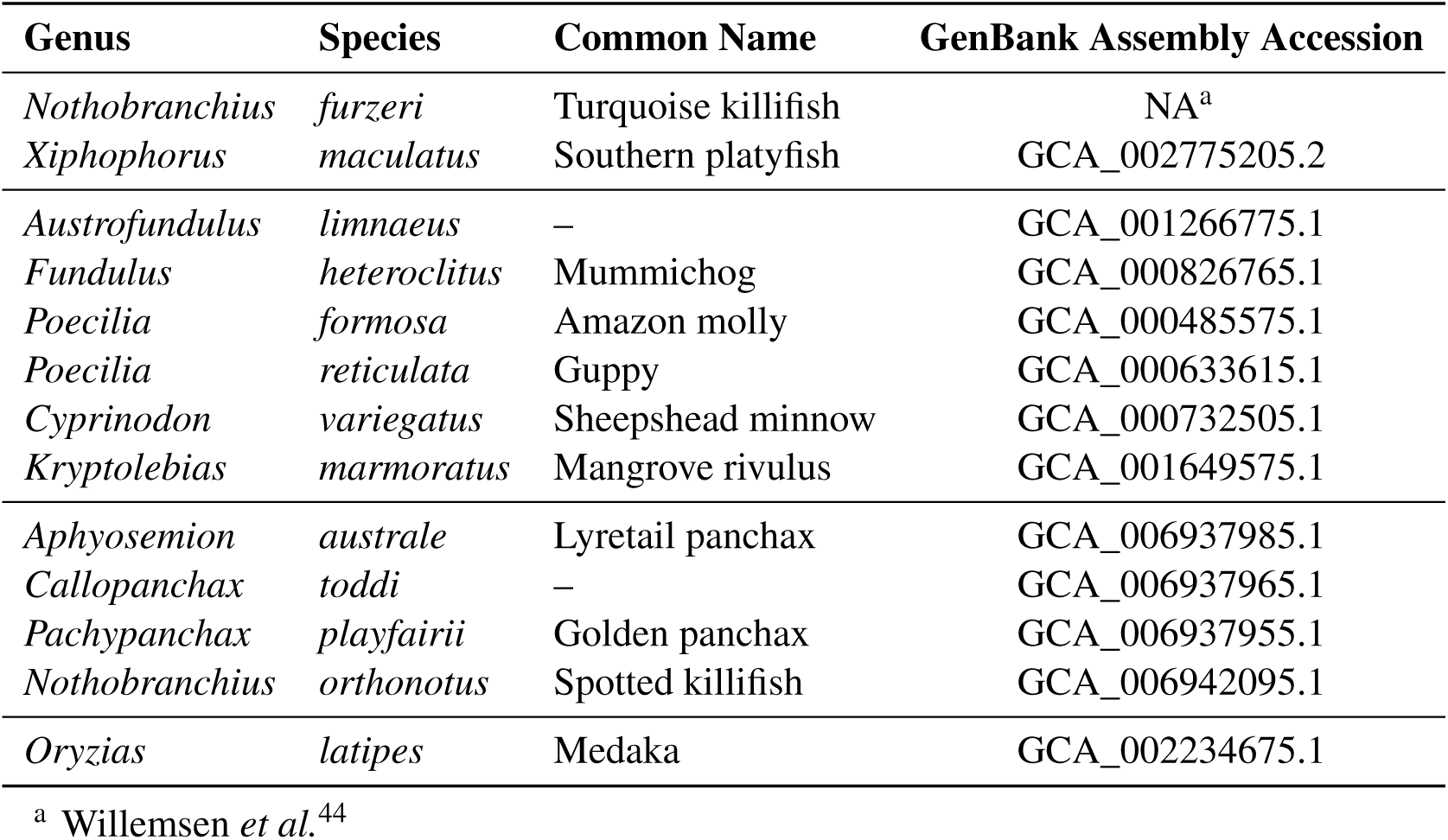
Genome assemblies used to identify *IGH* locus sequences in cyprinodontiform fishes

**Table S3:**
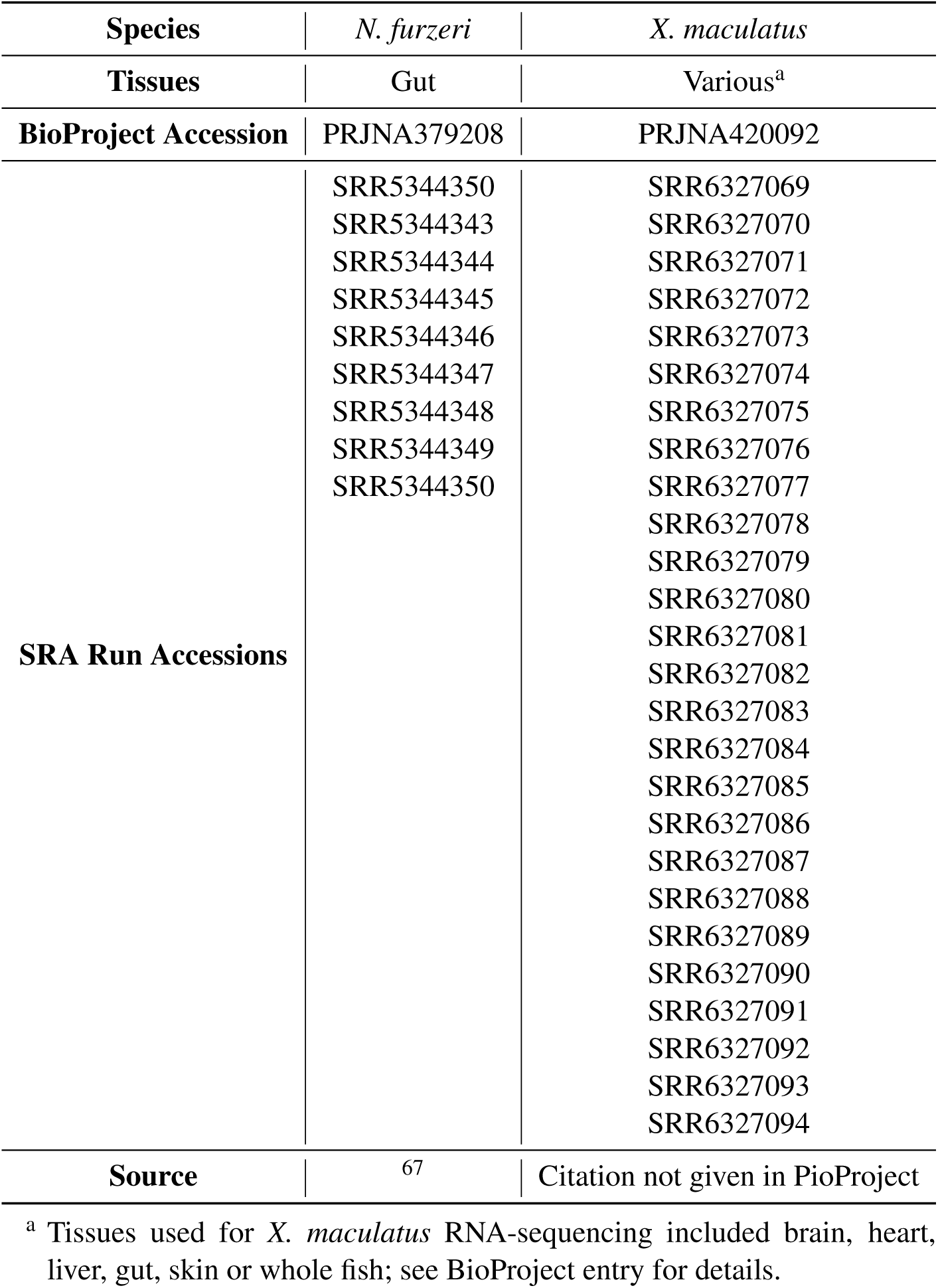
RNA-sequencing datasets used for *IGH* locus characterisation

**Table S4:** Co-ordinate table of constant-region exons in the *N. furzeri IGH* locus

| Name | Isotype | Start | End | Length | Strand |
| --- | --- | --- | --- | --- | --- |
| IGH1M-1 | M | 130848 | 131144 | 297 | + |
| IGH1M-2 | M | 131971 | 132312 | 342 | + |
| IGH1M-3 | M | 132394 | 132705 | 312 | + |
| IGH1M-4 | M | 132816 | 133288 | 473 | + |
| IGH1M-TM1 | M | 134262 | 134413 | 152 | + |
| IGH1M-TM2 | M | 138431 | 138819 | 389 | + |
| IGH1D-1 | D | 139381 | 139689 | 309 | + |
| IGH1D-2A | D | 139774 | 140064 | 291 | + |
| IGH1D-3A | D | 140178 | 140489 | 312 | + |
| IGH1D-4A | D | 140572 | 140853 | 282 | + |
| IGH1D-2B | D | 145613 | 145909 | 297 | + |
| IGH1D-3B | D | 146000 | 146311 | 312 | + |
| IGH1D-4B | D | 146398 | 146676 | 279 | + |
| IGH1D-5 | D | 146795 | 147124 | 330 | + |
| IGH1D-6 | D | 147210 | 147527 | 318 | + |
| IGH1D-7 | D | 147598 | 147885 | 288 | + |
| IGH1D-TM1 | D | 148016 | 148164 | 149 | + |
| IGH1D-TM2 | D | 148323 | 148504 | 182 | + |
| IGH2D-TM2 | D | 187624 | 187803 | 180 | - |
| IGH2D-TM1 | D | 187963 | 188111 | 149 | - |
| IGH2D-7 | D | 188658 | 188945 | 288 | - |
| IGH2D-6 | D | 189016 | 189333 | 318 | - |
| IGH2D-5 | D | 189419 | 189748 | 330 | - |
| IGH2D-4B | D | 189867 | 190145 | 279 | - |
| IGH2D-3B | D | 190232 | 190543 | 312 | - |
| IGH2D-2B | D | 190636 | 190932 | 297 | - |
| IGH2D-4A | D | 195644 | 195925 | 282 | - |
| IGH2D-3A | D | 196008 | 196319 | 312 | - |
| IGH2D-2A | D | 196433 | 196723 | 291 | - |
| IGH2D-1 | D | 196808 | 197116 | 309 | - |
| IGH2M-TM2 | M | 198315 | 198506 | 192 | - |
| IGH2M-TM1 | M | 199834 | 199985 | 152 | - |
| IGH2M-4 | M | 200953 | 201425 | 473 | - |
| IGH2M-3 | M | 201536 | 201847 | 312 | - |
| IGH2M-2 | M | 201929 | 202270 | 342 | - |
| IGH2M-1 | M | 203549 | 203845 | 297 | - |

**Table S5:**
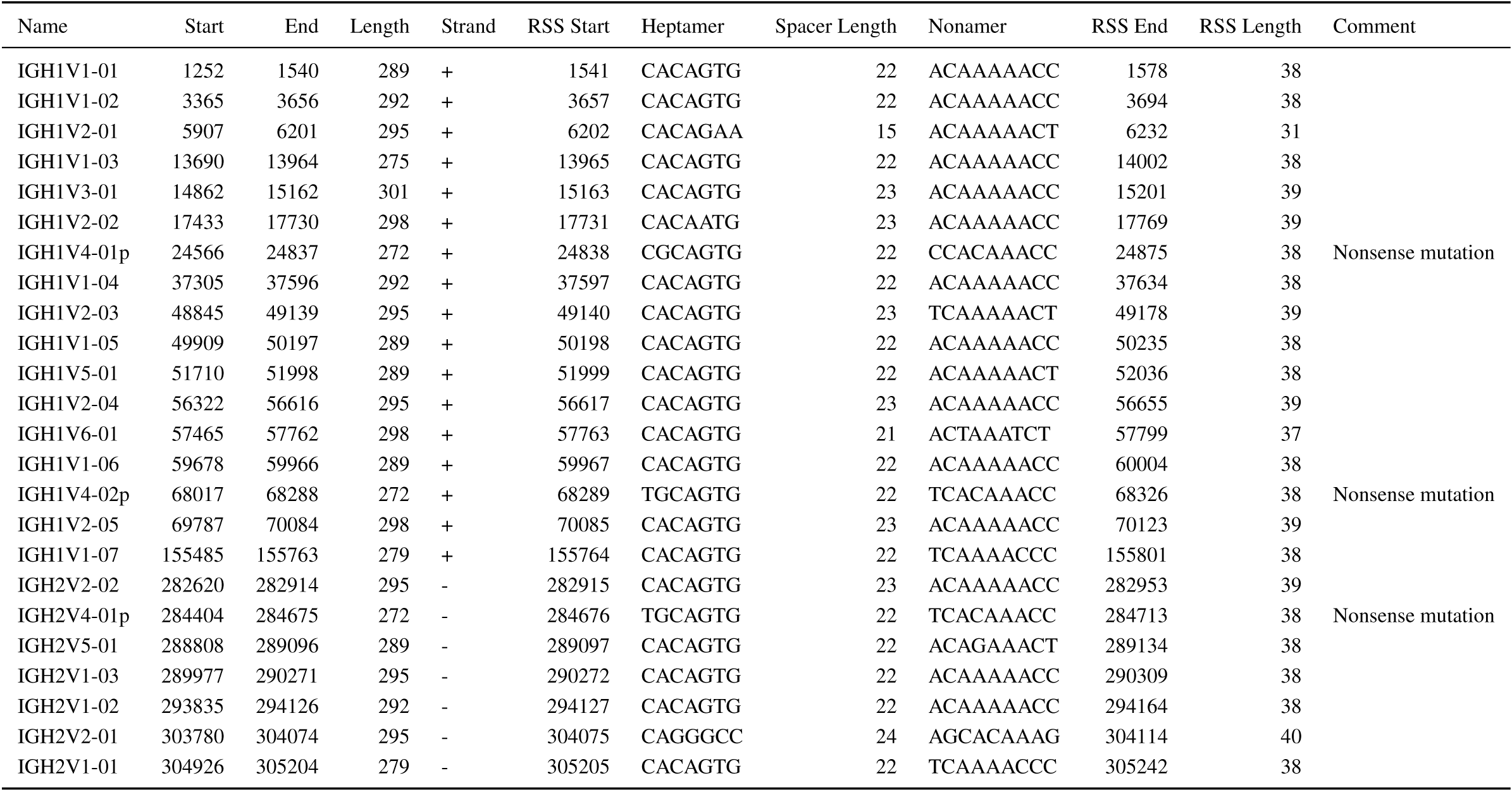
Co-ordinate table of VH segments in the *N. furzeri IGH* locus

**Table S6:**
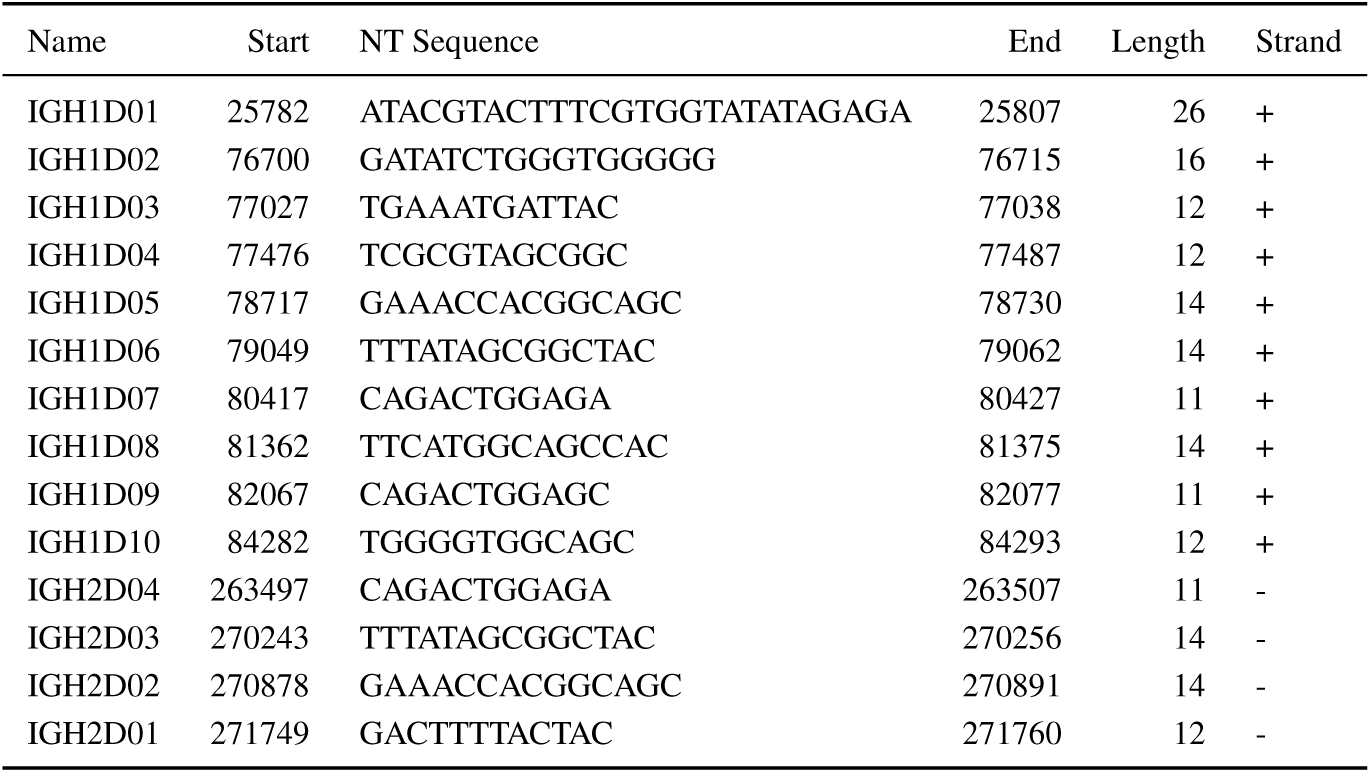
Co-ordinate table of DH segments in the *N. furzeri IGH* locus

**Table S7:**
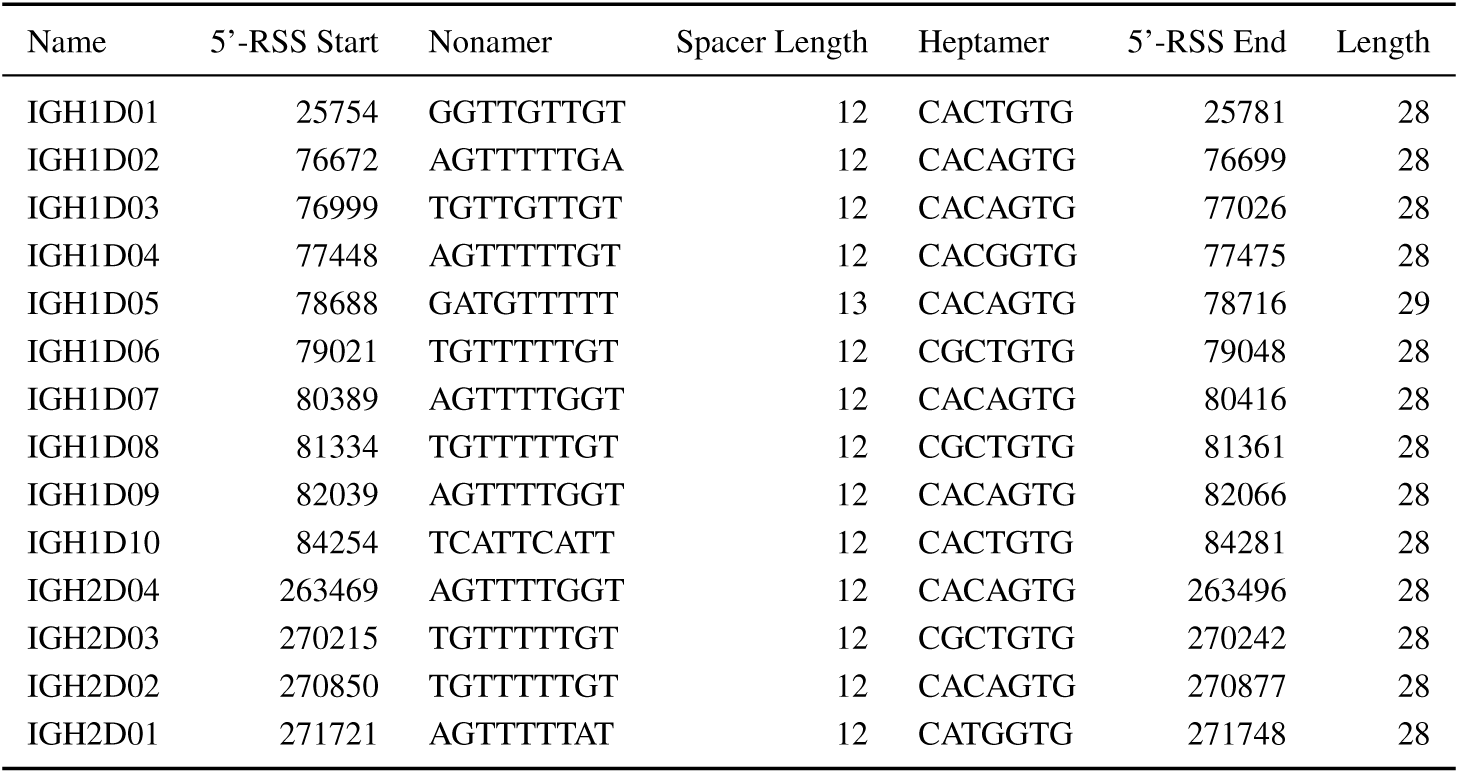
Co-ordinate table of DH 5’-RSSs in the *N. furzeri IGH* locus

**Table S8:**
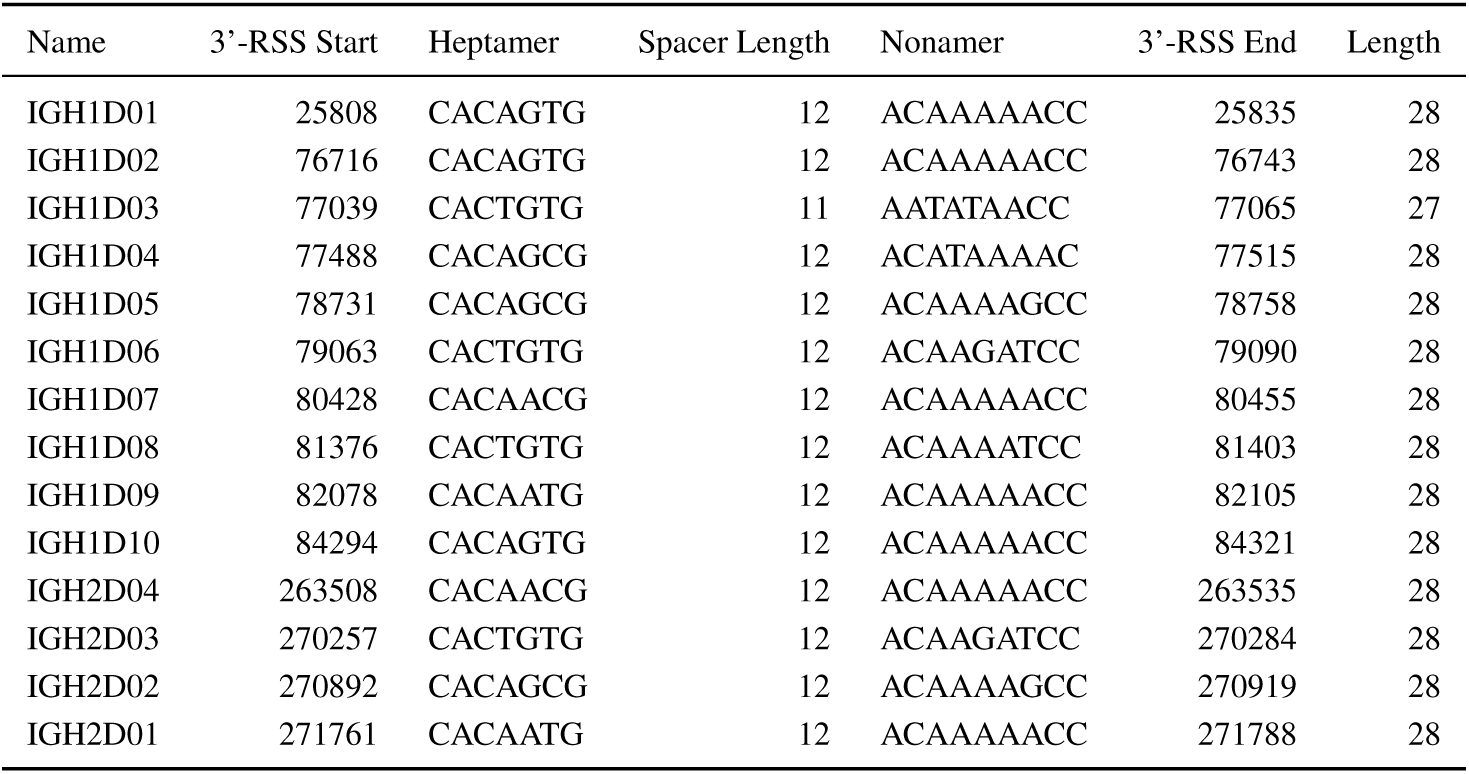
Co-ordinate table of DH 3’-RSSs in the *N. furzeri IGH* locus

**Table S9:**
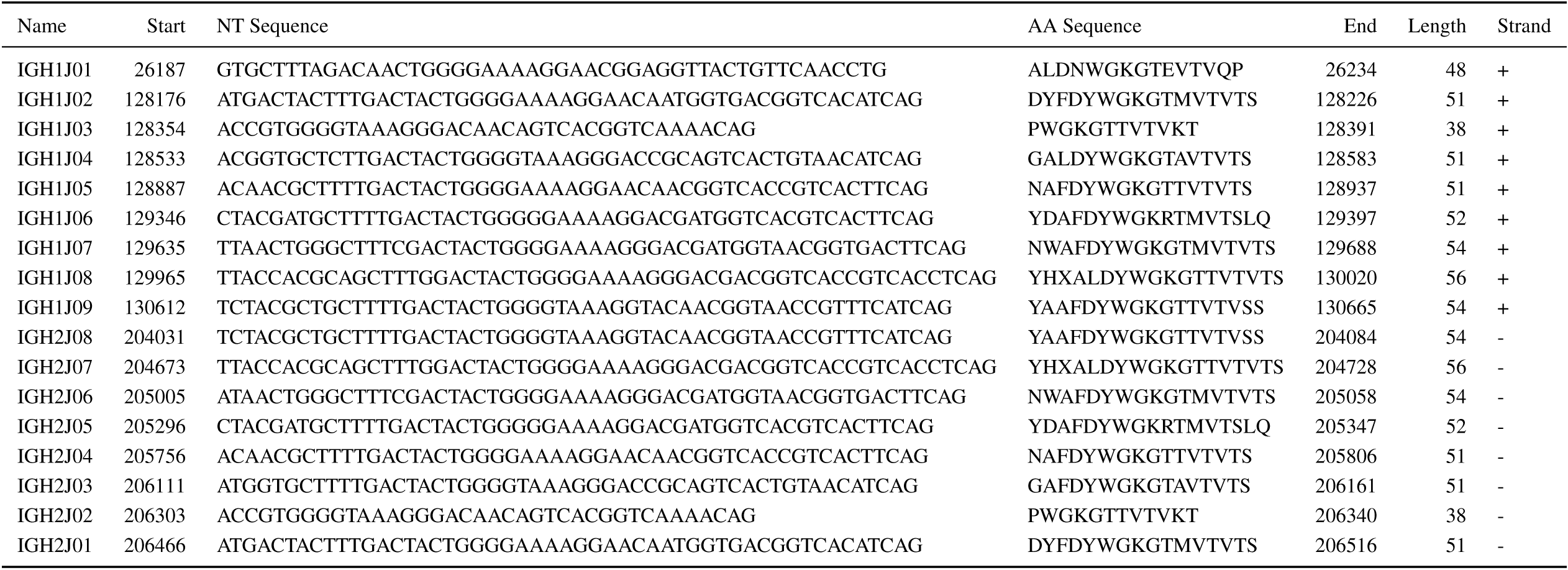
Co-ordinate table of JH segments in the *N. furzeri IGH* locus

**Table S10:**
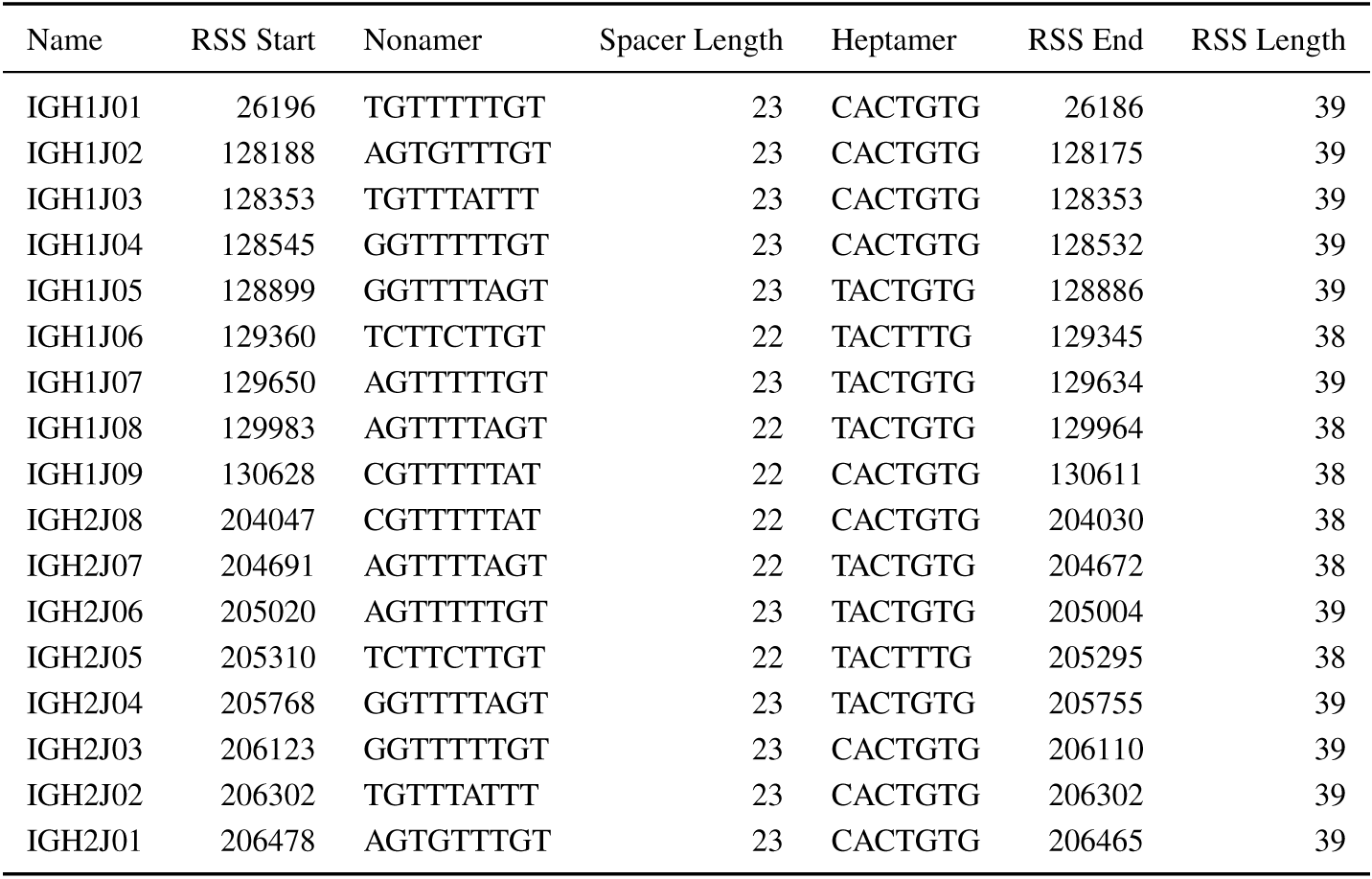
Co-ordinate table of JH RSSs in the *N. furzeri IGH* locus

**Table S11:** Co-ordinate table of constant-region exons in the *X. maculatus IGH* locus

| Name | Isotype | Start | End | Length | Strand |
| --- | --- | --- | --- | --- | --- |
| IGHZ1-1 | Z | 3380 | 3667 | 288 | + |
| IGHZ1-2 | Z | 3814 | 4098 | 285 | + |
| IGHZ1-3 | Z | 4195 | 4497 | 303 | + |
| IGHZ1-4 | Z | 4934 | 5263 | 330 | + |
| IGHZ1-S | Z | 5264 | 5459 | 196 | + |
| IGHZ1-TM1 | Z | 6345 | 6490 | 146 | + |
| IGHZ1-TM2 | Z | 6645 | 7043 | 399 | + |
| IGHZ2-1 | Z | 256059 | 256337 | 279 | + |
| IGHZ2-2 | Z | 256453 | 256734 | 282 | + |
| IGHZ2-3 | Z | 256893 | 257171 | 279 | + |
| IGHZ2-4 | Z | 257319 | 257636 | 318 | + |
| IGHZ2-S | Z | 257637 | 257850 | 214 | + |
| IGHZ2-TM1 | Z | 258059 | 258213 | 155 | + |
| IGHZ2-TM2 | Z | 258410 | 258629 | 220 | + |
| IGHM-1 | M | 279664 | 279960 | 297 | + |
| IGHM-2 | M | 280880 | 281224 | 345 | + |
| IGHM-3 | M | 281321 | 281629 | 309 | + |
| IGHM-4 | M | 281789 | 282291 | 503 | + |
| IGHM-TM1 | M | 282910 | 283034 | 125 | + |
| IGHM-TM2 | M | 285028 | 285740 | 713 | + |
| IGHD-1 | D | 285902 | 286219 | 318 | + |
| IGHD-2A | D | 286310 | 286597 | 288 | + |
| IGHD-3A | D | 286814 | 287128 | 315 | + |
| IGHD-4A | D | 287250 | 287534 | 285 | + |
| IGHD-2B | D | 288876 | 289166 | 291 | + |
| IGHD-3B | D | 289262 | 289576 | 315 | + |
| IGHD-4B | D | 289680 | 289964 | 285 | + |
| IGHD-5 | D | 290052 | 290381 | 330 | + |
| IGHD-6 | D | 290472 | 290789 | 318 | + |
| IGHD-7 | D | 290865 | 291152 | 288 | + |
| IGHD-TM1 | D | 291286 | 291434 | 149 | + |
| IGHD-TM2 | D | 291541 | 291642 | 102 | + |

**Table S12:**
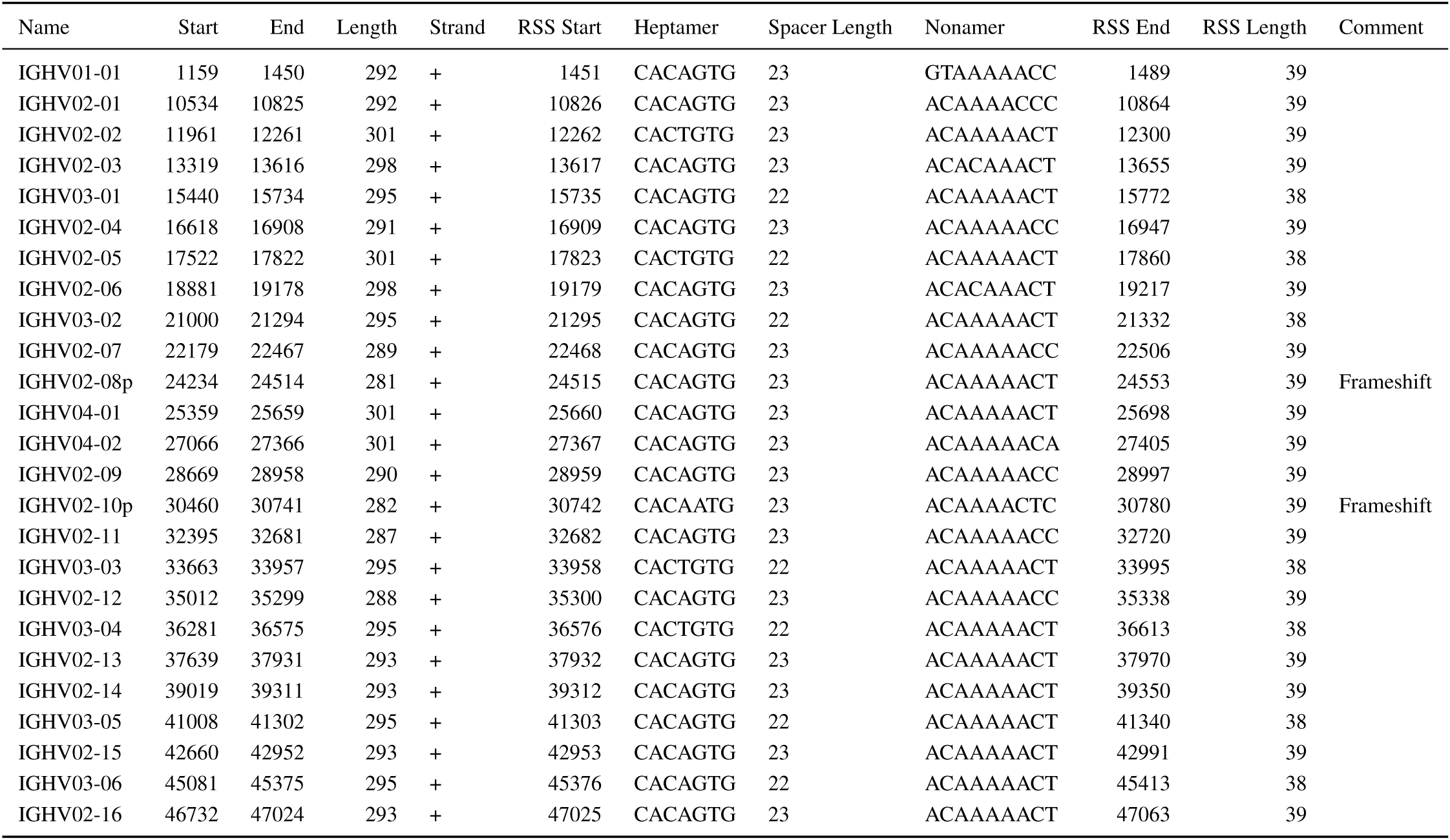
Co-ordinate table of VH segments in the *X. maculatus IGH* locus, part 1

**Table S13:**
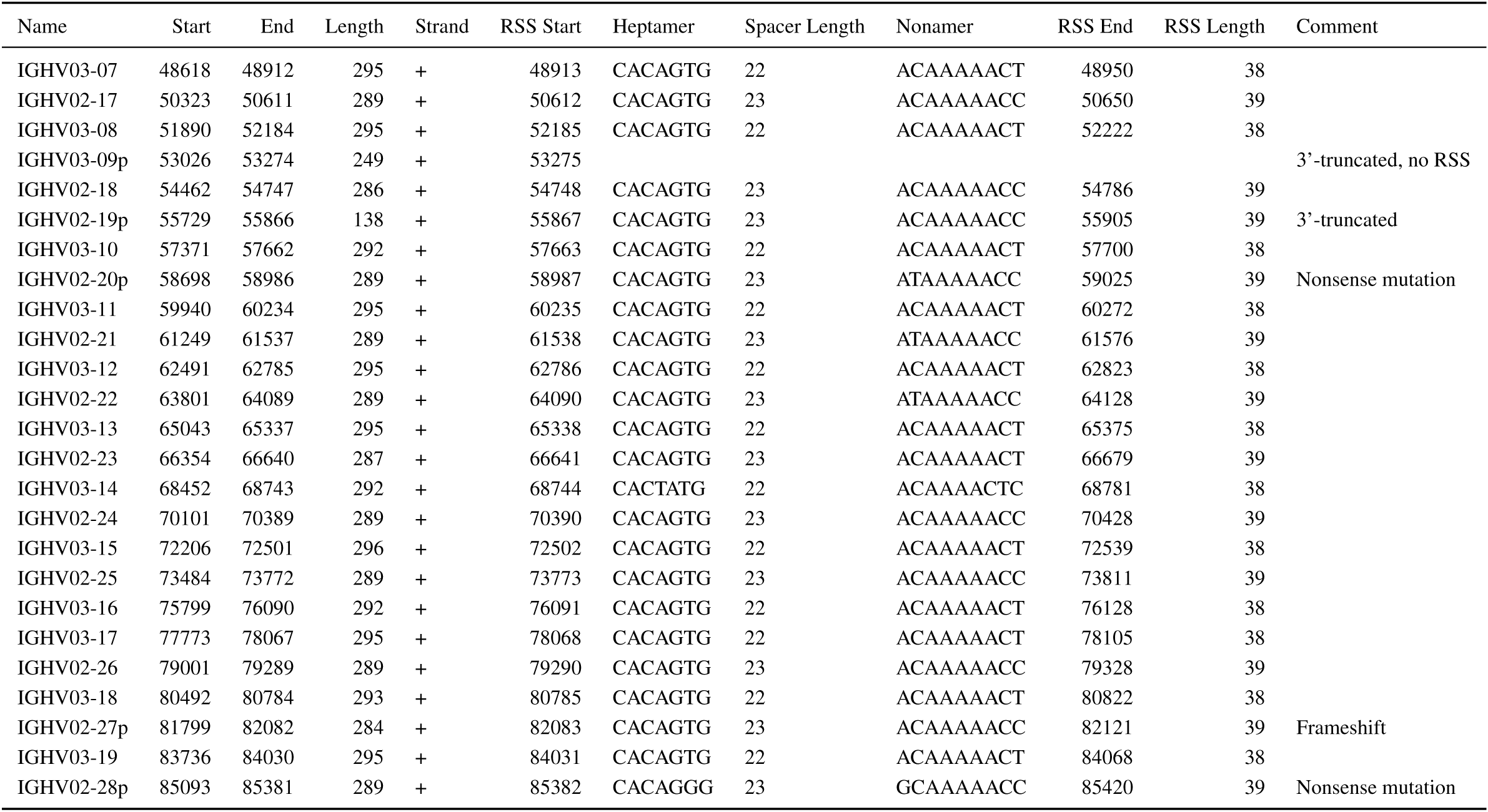
Co-ordinate table of VH segments in the *X. maculatus IGH* locus, part 2

**Table S14:**
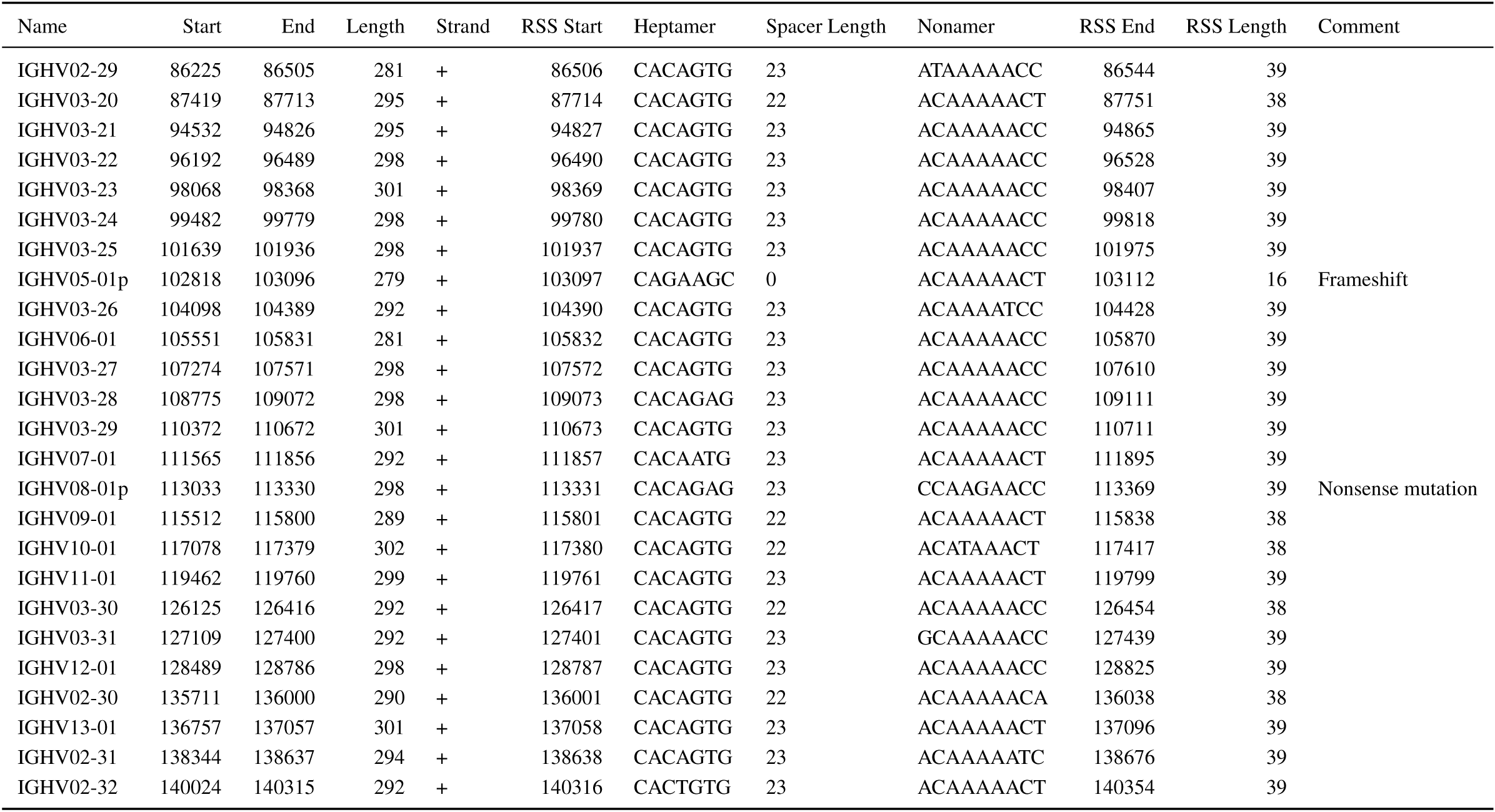
Co-ordinate table of VH segments in the *X. maculatus IGH* locus, part 3

**Table S15:**
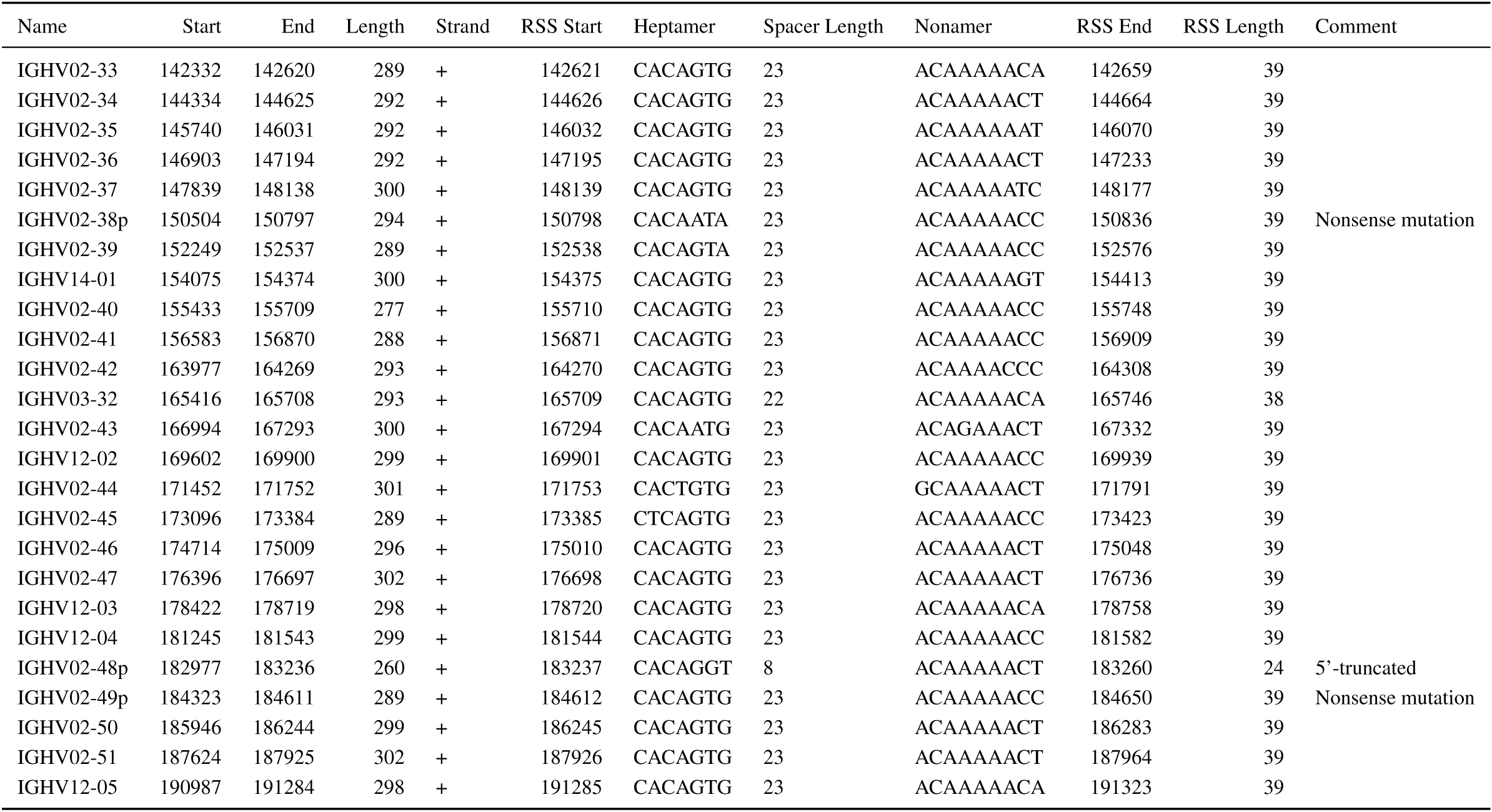
Co-ordinate table of VH segments in the *X. maculatus IGH* locus, part 4

**Table S16:**
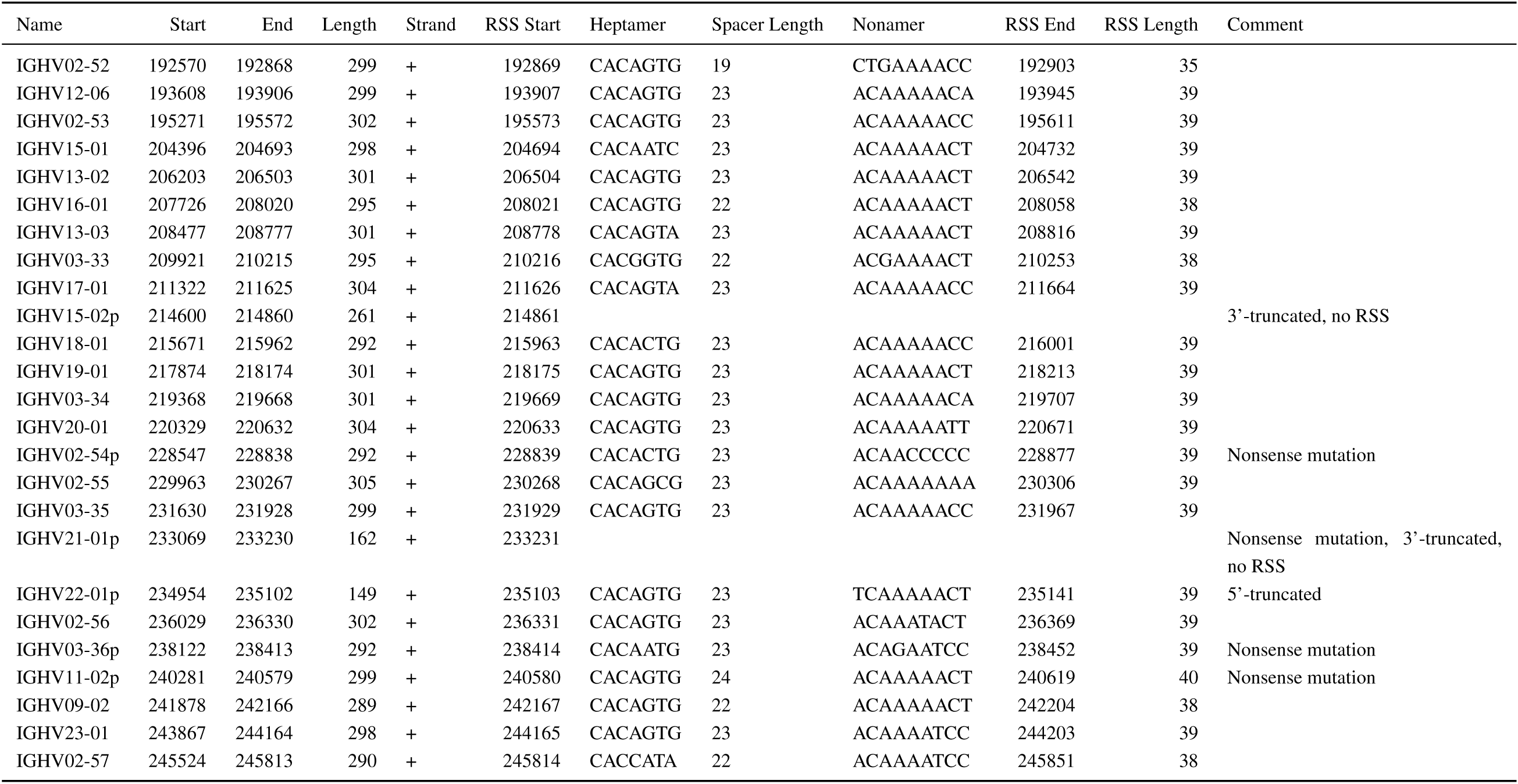
Co-ordinate table of VH segments in the *X. maculatus IGH* locus, part 5

**Table S17:**
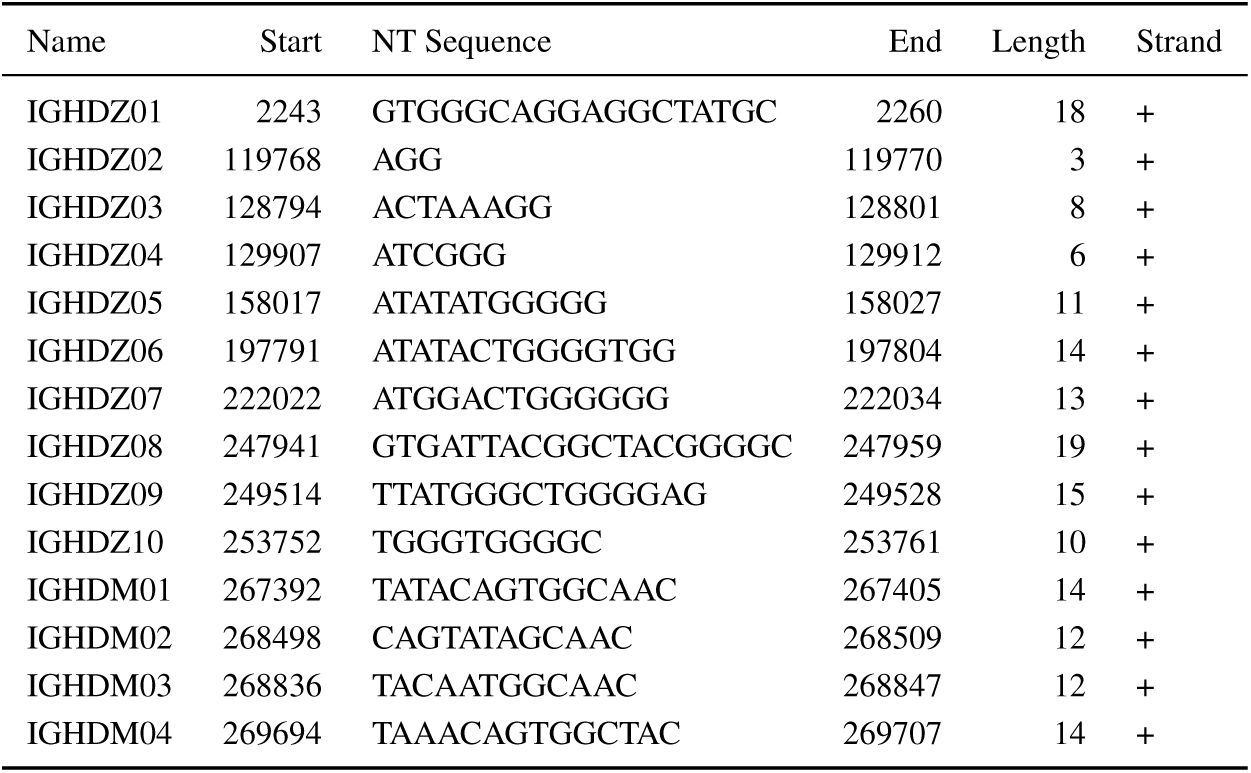
Co-ordinate table of DH segments in the *X. maculatus IGH* locus

**Table S18:** Co-ordinate table of DH 5’-RSSs in the *X. maculatus IGH* locus

| Name | 5'-RSS Start | Nonamer | Spacer Length | Heptamer | 5'-RSS End | Length |
| --- | --- | --- | --- | --- | --- | --- |
| IGHDZ01 | 2215 | GGTTTTTGT | 12 | CACTGTG | 2242 | 28 |
| IGHDZ02 | 119739 | TGTATTACT | 13 | CACAGTG | 119767 | 29 |
| IGHDZ03 | 128766 | TTTACTTCT | 12 | CACAGTG | 128793 | 28 |
| IGHDZ04 | 129879 | GGTTTTTGT | 12 | CACAGTG | 129906 | 28 |
| IGHDZ05 | 157989 | AGTTTTTGT | 12 | CACAGTG | 158016 | 28 |
| IGHDZ06 | 197763 | GGTTTTTGC | 12 | TACTGTG | 197790 | 28 |
| IGHDZ07 | 221994 | GGTTTTTGT | 12 | CGCTGTG | 222021 | 28 |
| IGHDZ08 | 247913 | TGTTTTTGT | 12 | ATCTGTG | 247940 | 28 |
| IGHDZ09 | 249486 | AGTTTTTGT | 12 | TGTGGTG | 249513 | 28 |
| IGHDZ10 | 253724 | AGTTTTTGT | 12 | TGTAGTG | 253751 | 28 |
| IGHDM01 | 267364 | AGTTTTTGT | 12 | TACAGTG | 267391 | 28 |
| IGHDM02 | 268470 | TGTTTTTGT | 12 | CACAGTG | 268497 | 28 |
| IGHDM03 | 268808 | AGTTTTTGC | 12 | TACTGTG | 268835 | 28 |
| IGHDM04 | 269666 | CGTTTTTGT | 12 | CATTGTG | 269693 | 28 |

**Table S19:**
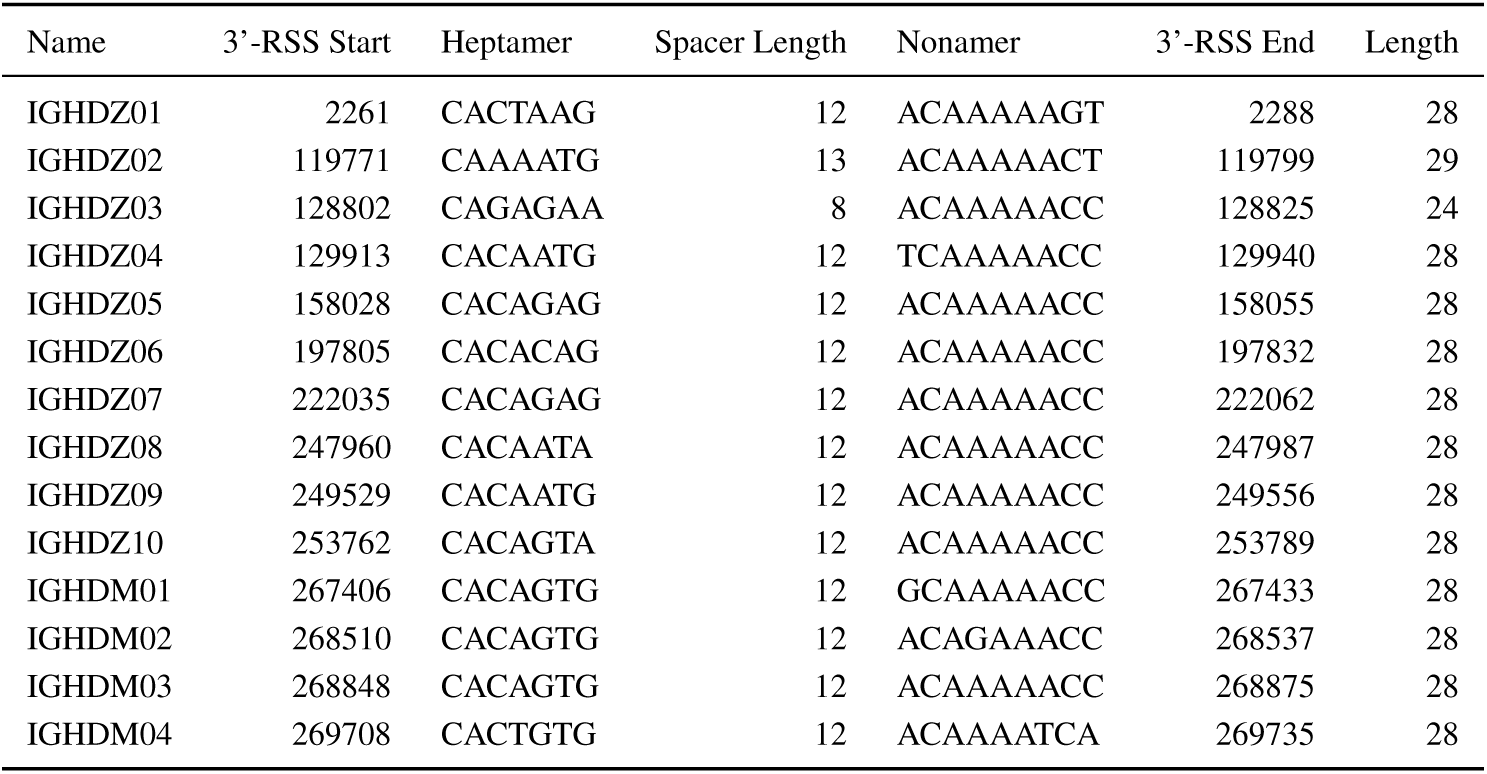
Co-ordinate table of DH 3’-RSSs in the *X. maculatus IGH* locus

**Table S20:**
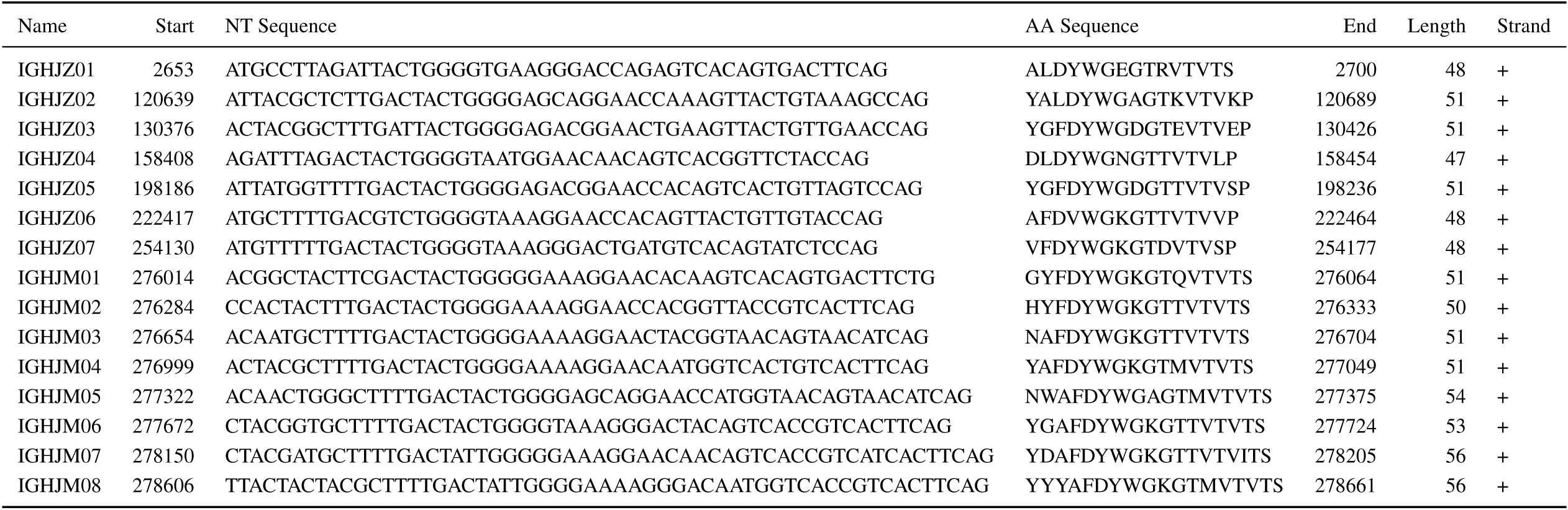
Co-ordinate table of JH segments in the *X. maculatus IGH* locus

**Table S21:** Co-ordinate table of JH RSSs in the *X. maculatus IGH* locus

| Name | RSS Start | Nonamer | Spacer Length | Heptamer | RSS End | RSS Length |
| --- | --- | --- | --- | --- | --- | --- |
| IGHJZ01 | 2662 | TGTTTTTGT | 23 | CACTGTG | 2652 | 39 |
| IGHJZ02 | 120651 | TGTTTTTGT | 23 | CACTGTG | 120638 | 39 |
| IGHJZ03 | 130388 | TGTTTTTGT | 23 | CACCGTG | 130375 | 39 |
| IGHJZ04 | 158416 | GGTTTTTGT | 23 | CACTGTG | 158407 | 39 |
| IGHJZ05 | 198198 | GGTTTTTGT | 23 | CACTGTG | 198185 | 39 |
| IGHJZ06 | 222426 | TGTTTTTGT | 23 | CACTGTG | 222416 | 39 |
| IGHJZ07 | 254139 | GGTTTTTGT | 23 | CACTGTG | 254129 | 39 |
| IGHJM01 | 276026 | TGTATTTGT | 23 | CACTGTG | 276013 | 39 |
| IGHJM02 | 276295 | TATTTTTGC | 23 | CACCGTG | 276283 | 39 |
| IGHJM03 | 276666 | TGTTTTTGT | 23 | TACTGTG | 276653 | 39 |
| IGHJM04 | 277011 | TGTTTTAGT | 23 | TACTGTG | 276998 | 39 |
| IGHJM05 | 277338 | GGTTTTTGT | 22 | TACTGTG | 277321 | 38 |
| IGHJM06 | 277687 | GCTTTTTAT | 22 | CACTGTG | 277671 | 38 |
| IGHJM07 | 278168 | CCTTTTTAC | 22 | CACTGTG | 278149 | 38 |
| IGHJM08 | 278624 | GCTTTTTAA | 22 | CACTGTG | 278605 | 38 |

**Table S22:**
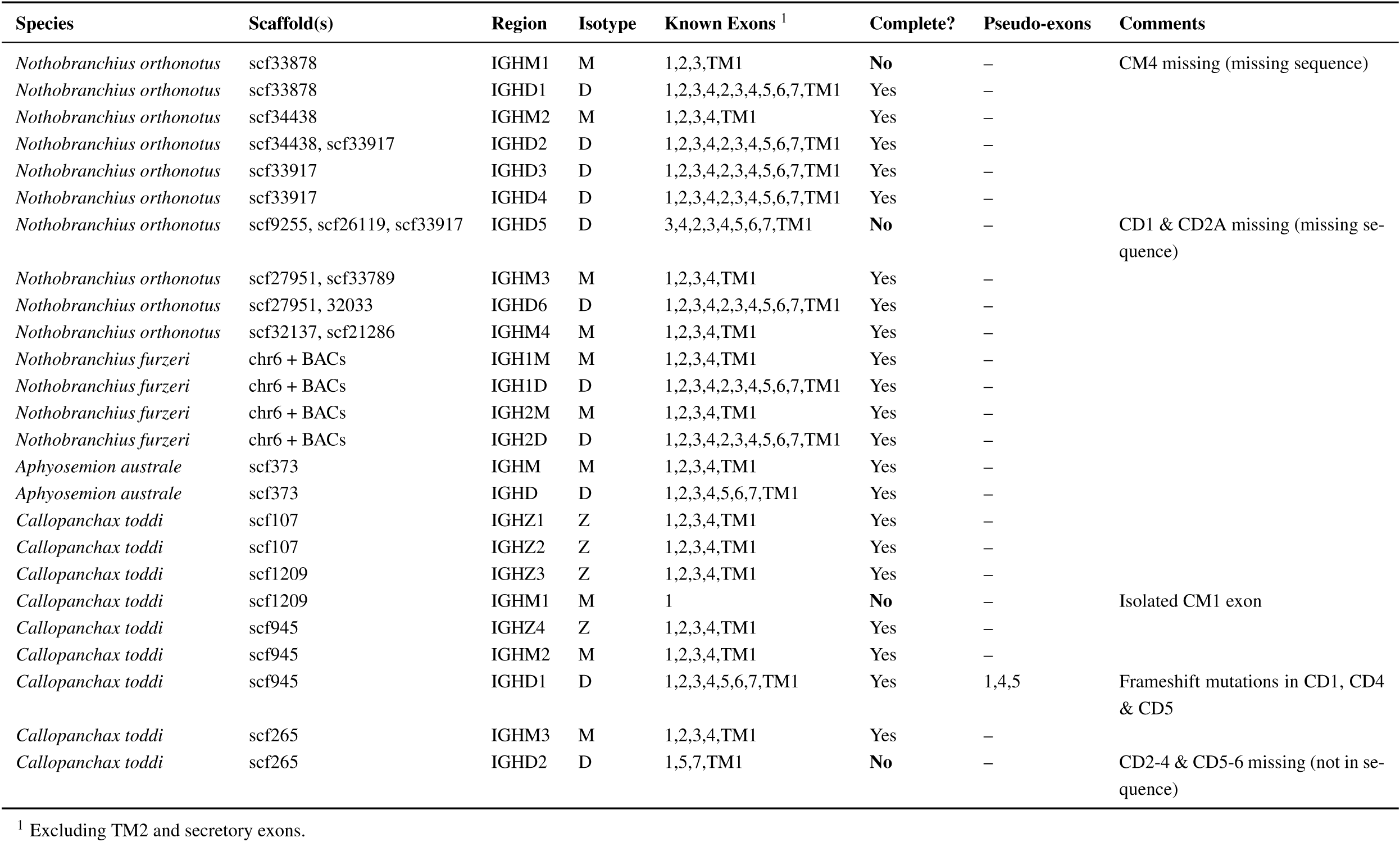
*IGH* constant regions in cyprinidontiform fish, part 1

**Table S23:** *IGH* constant regions in cyprinidontiform fish, part 2

| Species | Scaffold(s) | Region | Isotype | Known Exons <sup>1</sup> | Complete? | Pseudo-exons | Comments |
| --- | --- | --- | --- | --- | --- | --- | --- |
| *Pachypanchax playfairii* | scf547 | IGHZ | Z | 1,2,3,4,TM1 | Yes | - |  |
| *Pachypanchax playfairii* | scf125 | IGHM1 | M | 1,2,3,4,TM1 | Yes | - |  |
| *Pachypanchax playfairii* | scf125 | IGHD | D | 1,2,3,4,5,6,7,TM1 | Yes | - |  |
| *Pachypanchax playfairii* | scf547 | IGHM2 | M | 1 | No | - | Isolated CM1 exon |
| *Austrofundulus limnaeus* | NW_013954375.1 | IGHZ | Z | TM1 | No | TM1 | Isolated TM1 exon with frameshift mutation |
| *Austrofundulus limnaeus* | NW_013952673.1 | IGHM | M | 1,2,3,4,TM1 | Yes | - |  |
| *Austrofundulus limnaeus* | NW_013952673.1, NW_013956335.1 | IGHD | D | 1,2,3,4,5,6,7,TM1 | Yes | - |  |
| *Kryptolebias marmoratus* | NW_016094348.1 | IGHZ1 | Z | 1,2,3,4,TM1 | Yes | - |  |
| *Kryptolebias marmoratus* | NW_016094348.1 | IGHZ2 | Z | 1,4,TM1 | No | - | CZ2 & CZ3 missing (not in sequence) |
| *Kryptolebias marmoratus* | NW_016094301.1 | IGHM1 | M | 1,2,3,4,TM1 | Yes | - |  |
| *Kryptolebias marmoratus* | NW_016094301.1 | IGHD1 | D | 1,2,3,4,5,6,7,TM1 | Yes | - |  |
| *Kryptolebias marmoratus* | NW_016094277.1 | IGHM2 | M | 1,2,3,4,TM1 | Yes | - |  |
| *Kryptolebias marmoratus* | NW_016094277.1 | IGHD2 | D | 1,2,3,4,5,6,TM1 | No | - | CD7 missing (not in sequence) |
| *Poecilia reticulata* | NC_024338.1 | IGHZ1 | Z | 1,2,3,4 | No | - | TM1 missing (missing sequence) |
| *Poecilia reticulata* | NC_024338.1 | IGHZ2 | Z | 1,2,3,4,TM1 | Yes | - |  |
| *Poecilia reticulata* | NC_024338.1 | IGHM | M | 1,2,3,4,TM1 | Yes | - |  |
| *Poecilia reticulata* | NC_024338.1 | IGHD | D | 1,2,3,4,2,3,4,5,6,7,TM1 | Yes | - |  |
| *Poecilia formosa* | NW_006800081.1 | IGHZ1 | Z | 1,2,3,4,TM1 | Yes | - |  |
| *Poecilia formosa* | NW_006800081.1 | IGHZ2 | Z | 1,2,3,4,TM1 | Yes | - |  |
| *Poecilia formosa* | NW_006800081.1 | IGHZ3 | Z | 1,2,3,4,TM1 | Yes | - |  |
| *Poecilia formosa* | NW_006800081.1 | IGHM | M | 1,2,3,4,TM1 | Yes | - |  |
| *Poecilia formosa* | NW_006800081.1 | IGHD | D | 1,2,3,4,5,6,7,TM1 | Yes | - |  |
| *Xiphophorus maculatus* | NC_036458 | IGHZ1 | Z | 1,2,3,4,TM1 | Yes | - |  |
| *Xiphophorus maculatus* | NC_036458 | IGHZ2 | Z | 1,2,3,4,TM1 | Yes | - |  |
| *Xiphophorus maculatus* | NC_036458 | IGHM | M | 1,2,3,4,TM1 | Yes | - |  |

<sup>1</sup> Excluding TM2 and secretory exons.

**Table S24:** *IGH* constant regions in cyprinidontiform fish, part 3

| Species | Scaffold(s) | Region | Isotype | Known Exons <sup>1</sup> | Complete? | Pseudo-exons | Comments |
| --- | --- | --- | --- | --- | --- | --- | --- |
| *Xiphophorus maculatus* | NC_036458 | IGHD | D | 1,2,3,4,2,3,4,5,6,7,TM1 | Yes | – |  |
| *Fundulus heteroclitus* | NW_012234561.1 | IGHZ1 | Z | 1,2,3,4,TM1 | Yes | – |  |
| *Fundulus heteroclitus* | NW_012230737.1 | IGHZ2 | Z | 4,TM1 | No | – | CZ1 to CZ3 missing (missing sequence) |
| *Fundulus heteroclitus* | NW_012234542.1 | IGHM | M | 1,2,3,4,TM1 | Yes | – |  |
| *Fundulus heteroclitus* | NW_012234542.1 | IGHD | D | 1,2,3,4,2,3,4,5,6,7,TM1 | Yes | – |  |
| *Cyprinodon variegatus* | NW_015154250.1, NW_015151047.1 | IGHZ | Z | 1,2,3,4,TM1 | Yes | – |  |
| *Cyprinodon variegatus* | NW_015151047.1 | IGHM | M | 1,2,3,4,TM1 | Yes | – |  |
| *Cyprinodon variegatus* | NW_015151047.1 | IGHD | D | 1,2,3,4,2,3,4,5,6,7,TM1 | Yes | – |  |
| *Oryzias latipes* | NC_019866.2 | IGHM1 | M | 1,2,3,4,TM1 | Yes | – |  |
| *Oryzias latipes* | NC_019866.2 | IGHD1 | D | 1,2,3,4,6,7,TM1 | Yes | 7 | Nonsense mutation in CD7 |
| *Oryzias latipes* | NC_019866.2 | IGHM2 | M | 1,2,3,4,TM1 | Yes | – |  |
| *Oryzias latipes* | NC_019866.2 | IGHD2 | D | 1,2,3,4,6,7,TM1 | Yes | – |  |
| *Oryzias latipes* | NC_019866.2 | IGHM3 | M | 1,2,3,4,TM1 | Yes | – |  |
| *Oryzias latipes* | NC_019866.2 | IGHD3 | D | 1,2,3,4,6,7,TM1 | Yes | – |  |
| *Oryzias latipes* | NC_019866.2 | IGHM4 | M | 1,2,3,4,TM1 | Yes | – |  |
| *Oryzias latipes* | NC_019866.2 | IGHD4 | D | 2,7,TM1 | No | – | CD1 & CD3-6 missing (not in sequence) |
| *Oryzias latipes* | NC_019866.2 | IGHM5 | M | 1,2,3,4,TM1 | Yes | – |  |
| *Oryzias latipes* | NC_019866.2 | IGHD5 | D | 1,2,3,4,6,7,TM1 | Yes | – |  |
| *Oryzias latipes* | NC_019866.2 | IGHM6 | M | 1,2,3,4,TM1 | Yes | – |  |
| *Oryzias latipes* | NC_019866.2 | IGHD6 | D | 1,2,3,4,6,7,TM1 | Yes | – |  |
| *Oryzias latipes* | NC_019866.2 | IGHD7 | D | 1,2,3,6 | No | – | CD4, CD5, CD7 and TM1 missing (not in sequence) |

<sup>1</sup> Excluding TM2 and secretory exons.

## References

1. Jack, R. S. in Pathogen-Host Interactions: Antigenic Variation V. Somatic Adaptations (eds Hsu, E. & Du Pasquier, L.) 1–20 (Springer, 2015).

2. David Jung, Cosmas Giallourakis, Raul Mostoslavsky & Frederick W. Alt. Mechanism and Control of V(d)j Recombination at the Immunoglobulin Heavy Chain Locus. Annual Review of Immunology 24, 541–570 (2006).

3. Schatz, D. G. & Swanson, P. C. V(D)J Recombination: Mechanisms of Initiation. Annual Review of Genetics 45, 167–202 (2011).

4. Kurosaki, T., Kometani, K. & Ise, W. Memory B cells. Nature Reviews Immunology 15, 149–159 (2015).

5. Magor, B. G. Antibody Affinity Maturation in Fishes–Our Current Understanding. Biology 4, 512–524 (2015).

6. Mayer, A., Balasubramanian, V., Walczak, A. M. & Mora, T. How a well-adapting immune system re-members. arXiv, 1806.05753 (2018).

7. Schroeder, H. W. & Cavacini, L. Structure and function of immunoglobulins. Journal of Allergy and Clinical Immunology 125, S41–S52 (2010).

8. Mix, E., Goertsches, R. & Zett, U. K. Immunoglobulins–Basic considerations. Journal of Neurology 253, v9–v17 (2006).

9. Fillatreau, S. et al. The astonishing diversity of Ig classes and B cell repertoires in teleost fish. Frontiers in Immunology 4, 28 (2013).

10. Ravi, V. & Venkatesh, B. The divergent genomes of teleosts. Annual Review of Animal Biosciences 6, 47–68 (2018).

11. Danilova, N., Bussmann, J., Jekosch, K. & Steiner, L. A. The immunoglobulin heavy-chain locus in zebrafish: identification and expression of a previously unknown isotype, immunoglobulin Z. Nature Immunology 6, 295–302 (2005).

12. Magadán-Mompó, S., Sánchez-Espinel, C. & Gambón-Deza, F. Immunoglobulin heavy chains in medaka (*Oryzias latipes*). BMC Evolutionary Biology 11, 165 (2011).

13. Bao, Y. et al. The immunoglobulin gene loci in the teleost *Gasterosteus aculeatus*. Fish & Shellfish Immunology 28, 40–48 (2010).

14. Gambón-Deza, F., Sánchez-Espinel, C. & Magadán-Mompó, S. Presence of an unique IgT on the IGH locus in three-spined stickleback fish (*Gasterosteus aculeatus*) and the very recent generation of a repertoire of VH genes. Developmental & Comparative Immunology 34, 114–122 (2010).

15. Hansen, J. D., Landis, E. D. & Phillips, R. B. Discovery of a unique Ig heavy-chain isotype (IgT) in rainbow trout: Implications for a distinctive B cell developmental pathway in teleost fish. PNAS 102, 6919–6924 (2005).

16. Savan, R. et al. Discovery of a new class of immunoglobulin heavy chain from fugu. European Journal of Immunology 35, 3320–3331 (2005).

17. Yasuike, M. et al. Evolution of duplicated IgH loci in Atlantic salmon, *Salmo salar*. BMC Genomics 11, 486 (2010).

18. Bengtén, E. & Wilson, M. in Pathogen-Host Interactions: Antigenic Variation V. Somatic Adaptations (eds Hsu, E. & Du Pasquier, L.) 193–234 (Springer, 2015).

19. Hughes, L. C. et al. Comprehensive phylogeny of ray-finned fishes (Actinopterygii) based on transcriptomic and genomic data. PNAS 115, 6249–6254 (2018).

20. Valenzano, D. R. et al. The African Turquoise Killifish Genome Provides Insights into Evolution and Genetic Architecture of Lifespan. Cell 163, 1539–1554 (2015).

21. Reichwald, K. et al. Insights into Sex Chromosome Evolution and Aging from the Genome of a Short-Lived Fish. Cell 163, 1527–1538 (2015).

22. Cellerino, A., Valenzano, D. R. & Reichard, M. From the bush to the bench: the annual *Nothobranchius* fishes as a new model system in biology. Biological Reviews 91, 511–533 (2016).

23. Schartl, M. et al. The genome of the platyfish, *Xiphophorus maculatus*, provides insights into evolutionary adaptation and several complex traits. Nature Genetics 45, 567–572 (2013).

24. Altschul, S. F. et al. Basic local alignment search tool. Journal of Molecular Biology 215, 403–410 (1990).

25. Altschul, S. F. et al. Gapped BLAST and PSI-BLAST: a new generation of protein database search programs. Nucleic Acids Research 25, 3389–3402 (1997).

26. Bankevich, A. et al. SPAdes: A New Genome Assembly Algorithm and Its Applications to Single-Cell Sequencing. Journal of Computational Biology 19, 455–477 (2012).

27. Boetzer, M. et al. Scaffolding pre-assembled contigs using SSPACE. Bioinformatics 27, 578–579 (2011).

28. Sanger, F., Nicklen, S. & Coulson, A. R. DNA sequencing with chain-terminating inhibitors. PNAS 74, 5463–5467 (1977).

29. Bengtén, E. et al. Structure of the catfish IGH locus: analysis of the region including the single functional *IGHM* gene. Immunogenetics 58, 831–844 (2006).

30. Ramirez-Gomez, F. et al. Discovery and Characterization of Secretory IgD in Rainbow Trout: Secretory IgD Is Produced through a Novel Splicing Mechanism. The Journal of Immunology 188, 1341–1349 (2012).

31. Zhang, Y.-A. et al. IgT, a primitive immunoglobulin class specialized in mucosal immunity. Nature Immunology 11, 827–835 (2010).

32. Xu, Z. et al. Teleost skin, an ancient mucosal surface that elicits gut-like immune responses. PNAS 110, 13097–13102 (2013).

33. Mashoof, S. & Criscitiello, M. F. Fish Immunoglobulins. Biology 5, 45 (2016).

34. Dobin, A. et al. STAR: ultrafast universal RNA-seq aligner. Bioinformatics 29, 15–21 (2013).

35. Löytynoja, A. in Multiple Sequence Alignment Methods (ed Russell, D. J.) 155–170 (Humana Press, 2014).

36. Stamatakis, A. RAxML version 8: a tool for phylogenetic analysis and post-analysis of large phylogenies. Bioinformatics 30, 1312–1313 (2014).

37. Magurran, A. E. Evolutionary ecology: the Trinidadian guppy (Oxford University Press, Oxford, 2005).

38. Reid, N. M. et al. The genomic landscape of rapid repeated evolutionary adaptation to toxic pollution in wild fish. Science 354, 1305–1308 (2016).

39. Taylor, D. S. Twenty-four years in the mud: what have we learned about the natural history and ecology of the mangrove rivulus, *Kryptolebias marmoratus?* Integrative and Comparative Biology 52, 724–736 (2012).

40. Cui, R. et al. Relaxed selection limits lifespan by increasing mutation load. Cell 178, 1–15 (2019).

41. Weinstein, J. A., Jiang, N., White, R. A. & Quake, S. R. High-Throughput Sequencing of the Zebrafish Antibody Repertoire. Science 324, 807–810 (2009).

42. Georgiou, G. et al. The promise and challenge of high-throughput sequencing of the antibody repertoire. Nature Biotechnology 32, 158–168 (2014).

43. Jiang, N. et al. Determinism and stochasticity during maturation of the zebrafish antibody repertoire. PNAS 108, 5348–5353 (2011).

44. Willemsen, D., Cui, R., Reichard, M. & Valenzano, D. R. Genomics and population genetics in turquoise killifish reveal demography as driver for lifespan evolution (In preparation).

45. Bolger, A. M., Lohse, M. & Usadel, B. Trimmomatic: a flexible trimmer for Illumina sequence data. Bioinformatics 30, 2114–2120 (2014).

46. Langmead, B. & Salzberg, S. L. Fast gapped-read alignment with Bowtie 2. Nature Methods 9, 357–359 (2012).

47. Marçais, G., Yorke, J. A. & Zimin, A. QuorUM: An Error Corrector for Illumina Reads. PLOS One 10, e0130821 (2015).

48. Nikolenko, S. I., Korobeynikov, A. I. & Alekseyev, M. A. BayesHammer: Bayesian clustering for error correction in single-cell sequencing. BMC Genomics 14, S7 (2013).

49. Untergasser, A. et al. Primer3—new capabilities and interfaces. Nucleic Acids Research 40, e115 (2012).

50. Smith, A., Hubley, R. & Green, P. RepeatMasker Open-4.0. URL: https://www.repeatmasker.org (2018).

51. Li, H. et al. The Sequence Alignment/Map format and SAMtools. Bioinformatics 25, 2078–2079 (2009).

52. Robinson, J. T., et al. Integrative genomics viewer. Nature Biotechnology 29, 24–26 (2011).

53. Thorvaldsdóttir, H., Robinson, J. T. & Mesirov, J. P. Integrative Genomics Viewer (IGV): high-performance genomics data visualization and exploration. Briefings in Bioinformatics 14, 178–192 (2013).

54. Hahne, F. & Ivanek, R. in Statistical Genomics: Methods and Protocols (eds Mathé, E. & Davis, S.) 335–351 (Springer, 2016).

55. Shapiro, M. B. & Senapathy, P. RNA splice junctions of different classes of eukaryotes: sequence statistics and functional implications in gene expression. Nucleic Acids Research 15, 7155–7174 (1987).

56. Ulitsky, I. et al. Extensive alternative polyadenylation during zebrafish development. Genome Research 22, 2054–2066 (2012).

57. Wheeler, T. J. & Eddy, S. R. nhmmer: DNA homology search with profile HMMs. Bioinformatics 29, 2487–2489 (2013).

58. Hesse, J. E., Lieber, M. R., Mizuuchi, K. & Gellert, M. V(D)J recombination: a functional definition of the joining signals. Genes & Development 3, 1053–1061 (1989).

59. Ehrenmann, F. & Lefranc, M.-P. IMGT/DomainGapAlign: IMGT Standardized Analysis of Amino Acid Sequences of Variable, Constant, and Groove Domains (IG, TR, MH, IgSF, MhSF). Cold Spring Harbor Protocols 2011, 737–749 (2011).

60. Lefranc, M.-P. et al. IMGT unique numbering for immunoglobulin and T cell receptor variable domains and Ig superfamily V-like domains. Developmental & Comparative Immunology 27, 55–77 (2003).

61. Needleman, S. B. & Wunsch, C. D. A general method applicable to the search for similarities in the amino acid sequence of two proteins. Journal of Molecular Biology 48, 443–453 (1970).

62. Pagès, H., Aboyoun, P., Gentleman, R. & DebRoy, S. Biostrings: Efficient manipulation of biological strings. R package. URL: https://bioconductor.org/packages/release/bioc/html/biostrings.html (2019).

63. Rice, P., Longden, I. & Bleasby, A. EMBOSS: The European Molecular Biology Open Software Suite. Trends in Genetics 16, 276–277 (2000).

64. Yu, G., Lam, T. T.-Y., Zhu, H. & Guan, Y. Two methods for mapping and visualizing associated data on phylogeny using ggtree. Molecular Biology and Evolution 35, 3041–3043 (2018).

65. Rambaut, A. FigTree version 1.4. URL: https://tree.bio.ed.ac.uk/software/figtree/ (2018).

66. Wright, E. S. Using DECIPHER v2.0 to Analyze Big Biological Sequence Data in R. The R Journal 8, 352–359 (2016).

67. Smith, P. et al. Regulation of life span by the gut microbiota in the short-lived African turquoise killifish. eLife 6, e27014 (2017).

68. Pathogen-Host Interactions: Antigenic Variation V. Somatic Adaptations (eds Hsu, E. & Du Pasquier, L.) (Springer, 2015).

